# Loss of GABA co-transmission from cholinergic neurons impairs behaviors related to hippocampal, striatal, and medial prefrontal cortex functions

**DOI:** 10.1101/2022.10.07.511349

**Authors:** R. Oliver Goral, Kathryn M. Harper, Briana J. Bernstein, Sydney A. Fry, Patricia W. Lamb, Sheryl S. Moy, Jesse D. Cushman, Jerrel L. Yakel

## Abstract

Altered signaling or function of acetylcholine (ACh) has been reported in various neurological diseases, including Alzheimer’s disease, Tourette syndrome, epilepsy among others. Many neurons that release ACh also co-transmit the neurotransmitter gamma-aminobutyrate (GABA) at synapses in the hippocampus, striatum, and medial prefrontal cortex (mPFC). Although ACh transmission is crucial for higher brain functions such as learning and memory, the role of co-transmitted GABA from ACh neurons in brain function remains unknown. Thus, the overarching goal of this study was to investigate how a systemic loss of GABA co-transmission from ACh neurons affected the behavioral performance of mice. To do this, we used a conditional knock-out mouse of the vesicular GABA transporter (vGAT) crossed with the ChAT-Cre driver line to selectively ablate GABA co-transmission at ACh synapses. In a comprehensive series of standardized behavioral assays, we compared Cre-negative control mice with Cre-positive vGAT knock-out mice of both sexes. Loss of GABA co-transmission from ACh neurons did not disrupt the animal’s sociability, motor skills or sensation. However, in the absence of GABA co-transmission, we found significant alterations in social, spatial and fear memory as well as a reduced reliance on striatum-dependent response strategies in a T-maze. In addition, male CKO mice showed increased locomotion. Taken together, the loss of GABA co-transmission leads to deficits in higher brain functions and behaviors. Therefore, we propose that ACh/GABA co-transmission modulates neural circuitry involved in the affected behaviors.

## Introduction

In agreement with Dale’s principle, most synaptic vesicles (SVs) contain only a single neurotransmitter type defined by the vesicular transporters located within its membrane (Gutierrez (2009)). In addition to neurotransmitters, SVs co-release protons, ATP and, in some cases, Zn^2+^ ions (Li et al. (2020), Burnstock (1976), Ellis and Burnstock (1989), Soto et al. (2018), Upmanyu et al. (2022)). Rarely, different neurotransmitters or neuropeptides are co-released from the same SVs (Tritsch et al. (2016), Wojcik et al. (2006)). More often, separate pools of SVs with different neurotransmitters and different intrinsic properties are released from the same neuron or synapse during a process called co-transmission (Takacs et al. (2018)).

Acetylcholine- (ACh) releasing neurons exist both as local interneurons as well as projection neurons and they form complex axonal arborizations with vast numbers of synapses which transmit signals through ionotropic nicotinic ACh receptors (nAChR) or metabotropic muscarinic ACh receptors (mAChR) (Ballinger et al. (2016)). The expression of nAChR and mAChR depends on pre- and/or postsynaptic location, cell type, cell location, and brain region (Guillem et al. (2011), Poorthuis et al. (2013)). Furthermore, effects of ACh transmission strongly depend on the timing of the ACh signal and other firing events (Gu and Yakel (2011), Unal et al. (2015)). Individual ACh neurons form synapses with many target neurons, even within different brain regions (Li et al. (2018), Nelson and Mooney (2016)). This enables the cholinergic system to coordinate the activity within a large network of neurons in the brain. In addition to modulating neuron firing properties, SV release probability and morphology, ACh signals can directly trigger the release of neuromodulators such as dopamine (DA) from axonal varicosities (Picciotto et al. (2012), Cheng and Yakel (2015), Urban-Ciecko et al. (2018), Morley and Mervis (2013), Lozada et al. (2012), Steinecke et al. (2022), Shin et al. (2017), Liu et al. (2022)). The varied effects of ACh transmission are required to coordinate oscillatory events of computational processes within the hippocampus or cortical areas where large neuron populations fire synchronously (Gu et al. (2017), Gu et al. (2020), Hasselmo and McGaughy (2004), Marrosu et al. (1995)).

The role of ACh release in cognition is well established (Ballinger et al. (2016)). It is, however, unclear why large subpopulations of ACh neurons co-express markers (e. g. GAD1/2, vGAT, Lhx6, VIP), which are typically found in gamma-amino-butyrate (GABA)-ergic neurons (Lee et al. (2010), Saunders et al. (2015a), Saunders et al. (2015b), Sethuramanujam et al. (2016), Takacs et al. (2018), Lozovaya et al. (2018), Obermayer et al. (2019), Granger et al. (2020)). These GABAergic ACh neurons can release GABA at their synapses in the retina, hippocampus, striatum, and medial prefrontal cortex (mPFC) (Lee et al. (2010), Saunders et al. (2015a), Saunders et al. (2015b), Sethuramanujam et al. (2016), Takacs et al. (2018) Lozovaya et al. (2018), Obermayer et al. (2019), Granger et al. (2020), Hunt et al. (2022)). There are indications that both neurotransmitters are co-transmitted at the same ACh synapses by distinct mechanisms (Lee et al. (2010), Takacs et al. (2018)). In fact, GABA is released with faster kinetics, requires less calcium, and relies mostly on Ca_V_2.1 channels indicating tight SV coupling; ACh release, in contrast, occurs with loose SV coupling (Neher and Sakaba (2008), Lee et al. (2010), Takacs et al. (2018)).

Co-transmission of ACh and GABA has been implicated in different aspects of striatal, hippocampal, and cortical function. In the striatum, ACh/GABA interneurons (CGINs) are more strongly involved in the pause response and more sensitive to local inhibition (Lozovaya (2018)). In a model of Parkinson’s disease (PD), co-transmitted excitatory GABA enlarged the dendritic fields of CGINs, causing an increased CGIN-CGIN connectivity and firing (Lozovaya (2018)). In the hippocampus, both ACh and GABA were found to be released at the same synapses onto oriens lacunosum moleculare (OLM) interneurons (Takacs (2018)). The release of GABA was sufficient to suppress sharp-wave ripples and epileptiform activity within the hippocampus (Takacs (2018)). The mPFC receives ACh/GABA inputs mostly onto L1 interneurons from the external segment of the globus pallidus (GPe) and from the adjacent part of the nucleus basalis of Meynert (NB) (Saunders et al. (2015a), Saunders et al. (2015b)). Moreover, VIP^+^/ChAT^+^ neurons intrinsic to the mPFC preferentially co-transmit ACh/GABA onto L1 interneurons with differences between model organisms (Obermayer et al. (2019), Granger et al. (2020)). Silencing mPFC VIP^+^/ChAT^+^ neurons negatively affected animal attention (Obermayer et al. (2019)). Furthermore, GABA co-transmission may modulate spike timing in target neurons (Obermayer et al. (2019)).

However, little is known about the importance of GABA co-transmission from ACh neurons for animal behavior. Therefore, we assessed whether mice without GABA co-transmission from ACh neurons showed any behavioral deficits. We ablated GABAergic transmission from ACh neurons by knocking out the vesicular GABA transporter (vGAT, Slc32a1) specifically in ACh neurons, and then evaluated the conditional knockout (CKO) mice in a battery of standardized behavior assays. We found that vGAT CKO mice had impaired social and spatial memory as well as minor alterations in striatal response learning and fear renewal. Furthermore, male CKO mice had increased locomotor activity. Taken together, these alterations indicate the involvement of GABA co-transmission from ACh neurons in complex behaviors that require hippocampal, striatal, and mPFC circuitry.

## Material and Methods

### Animals

Transgenic animal lines were purchased from The Jackson Laboratories and subsequently maintained and bred in-house. Offspring were group housed (<5 per cage, separated by sex) whenever possible in a regular 12 h light/dark cycle. Food and water were supplied *ad libitum*. To delete GABAergic co-transmission from cholinergic neurons, we crossed vGAT-flox mice (Jax# 012897, RRID:IMSR_JAX: 012897), with ChAT-IRES-Cre mice (Jax# 006410, RRID:IMSR_JAX:006410) (Tong et al. (2008), Rossi et al. (2011)). Mice were bred with homozygous vGAT-flox alleles, heterozygous ChAT-IRES-Cre alleles to obtain Cre-negative (ctrl) as well as heterozygous Cre-positive littermates (CKO) for behavior experiments.

All animal care and procedures were conducted in strict compliance with the animal welfare policies set by the National Institutes of Health. All procedures were approved and performed in compliance with the NIEHS/NIH Humane Care and Use of Animals Protocols, and, where applicable, by the UNC Institutional Animal Care and Use Committee and by the University of North Carolina at Chapel Hill (UNC).

### Animal Behavior Experiments

For general behavioral phenotyping, the UNC Behavioral Phenotyping Laboratory used 11 male and 11 female ctrl mice as well as 11 male and 13 female CKO mice. One male CKO mouse was withdrawn from the study before the start of water maze testing due to fighting wounds. Mice were 6-8 weeks of age at the beginning of the behavioral assessment. See Table 1 for detailed timeline.

**Table 1:**

| Age (weeks) | Procedure |
| --- | --- |
| 6-8 | Elevated plus maze for anxiety-like behavior |
| 6-9 | Locomotor activity and exploration in a 1-hour open field test |
| 7-10 | Rotarod test for motor coordination |
| 8-11 | Social approach in a three-chamber choice test |
| 9-12 | Marble-bury assay for anxiety-like behavior and perseverative responses<br>Acoustic startle test for pre-pulse inhibition |
| 10-13 | Buried food test for olfactory function |
| 11-16 | Visible platform test in Morris water maze |
| 12-17 | Hidden platform test for spatial learning in Morris water maze |
| 13-18 | Reversal learning test for cognitive flexibility in Morris water maze |
| 14-20 | Conditioned fear test for contextual and cue-dependent learning |

The T maze was performed at the NIEHS animal facility with a total of 64 mice (15 male ctrl, 15 male CKO, 18 female ctrl, 16 female CKO). Mice were 12-16 weeks of age at the beginning of the behavioral testing. The mice were placed in a 12 h reverse-light cycle room and put on a food-restriction schedule to gradually reduce and maintain the body weight at >85%. During a period of 7-10 days, the animals were weighed and handled (10 min/day) daily. Before the start of the behavioral testing, animals were acclimated to the room for >30 min. Behavioral testing was conducted during the animals’ dark cycle.

A total of 40 mice (8 male and 11 female ctrl mice and 8 male and 13 female CKO mice) was used for additional behavioral tests: 1) automated home-cage discrimination and reversal learning test, and 2) fear conditioning/extinction/renewal. One male CKO and one female ctrl mouse were excluded from the CognitionWall experiment after day 2 due to not reaching the 100-sugar pellet criterion. One male ctrl mouse died before the start of the fear conditioning/extinction/renewal experiments. Mice were 8–12 weeks in age at the beginning of behavioral testing. Both tasks were separated by about 4 weeks.

### Three-chamber choice test

The procedure was comprised of three 10-min phases: habituation, sociability test, and social novelty preference test. During the sociability test, mice had a choice between proximity to an unfamiliar, sex matched C57BL/6J adult mouse (“stranger 1”) and being alone. During the social novelty test, mice were to choose between the already-investigated stranger 1, and a new unfamiliar mouse (“stranger 2”). A rectangular, 3-chambered clear Plexiglas box with dividing walls and doorways to each chamber was used to perform the test. An automated image tracking system (Ethovision 7, Noldus, RRID:SCR_000441) provided measures of time spent in each side and entries into each side of the social test box.

During the first phase of the test, the mouse was placed in the middle chamber and allowed to explore for 10 minutes, with the doorways open into both side chambers. After the habituation phase, the test mouse was enclosed in the center compartment of the social test box, and stranger 1 was placed in one of the side chambers. The stranger mouse was enclosed in a small Plexiglas cage drilled with holes to allow nose contact. An identical empty Plexiglas cage was placed in the opposite chamber. After placement of stranger 1 and the empty cage, the doors were re-opened, and the subject was allowed to explore the social test box for another 10-min session. At the end of the sociability phase, stranger 2 was placed in the empty Plexiglas cage, and the subject was given an additional 10 minutes to explore the social test box.

### Morris water maze

Spatial and reversal learning, swimming ability, and vision were assessed by performance in the Morris water maze. A large circular pool (122 cm diameter) was partially filled with water (45 cm depth, 24-26°C), located in a room with numerous visual cues. The test consisted of three different phases: the visible platform test, acquisition in the hidden platform test, and the reversal learning test.

During the visible platform test, every mouse performed four trials per day on two consecutive days, to swim to an escape platform cued by a patterned cylinder extending above the water. The mouse started each trial in the pool at 1 of 4 possible locations at random and was given 60 s to find the visible platform. If the mouse reached the platform, the trial ended, and the animal remained on the platform for 10 s before the next trial began. In case the platform was not found, the mouse was placed on the platform for 10 s, and then given the next trial. Latency to reach the platform and swimming speed were measured via an automated tracking system (Ethovision 15).

Following the visible platform test, mice were tested for ability to find a submerged, hidden escape platform (diameter = 12 cm). Each mouse performed 4x 1 min trials per day, to swim to the hidden platform. In the present study, all groups reached the learning criterion on day 4 (15 s or less to locate the platform) and were given a 1-min probe trial in the pool without any platform on the same day. The selective search for the correct location was assessed by comparing number of swim path crosses over the quadrant where the platform (target) was positioned during training with the corresponding area in the opposite quadrant.

Following the hidden platform phase, mice were tested for reversal learning. During this phase, the same procedure was used as described above but with the hidden platform transferred to the opposite quadrant in the pool. As during the visible platform test, the latency to find the platform was assessed. On day 4 of testing, the platform was removed from the pool, and every mouse performed a probe trial to evaluate reversal learning. Quadrant preference was assessed by comparing swim paths over target and opposite quadrant.

### T maze

A plus maze with a white polypropylene floor (Med Associates) with four identical arms (37.47 × 8.92 × 14.68 cm) was raised to 17.46 cm and placed in a black-curtained area. All trials were recorded on video and tracked with Ethovision 15. Animals were trained during their dark cycle in the presence of dim white illumination with large white fabric cues (triangle, oval, rectangle, square) affixed to the curtain behind every arm. During habituation and training trials, the maze arm facing the start arm was blocked to create a T maze. Before the training, every mouse performed two 5 min habituation trials on each of two consecutive days to adjust to the maze environment. To avoid a direction bias, about half of the animals were trained to turn right to gain the reward of two sugar pellets (Dustless Precision Rodent Pellets, F05684, Bio-Serv, Flemington, NJ) located at the end of the right arm. The other half were trained to turn left for the reward. On each trial, the mouse was placed at the end of the start arm facing the center of the maze and given 2 min to find and eat the reward. After 2 min or when the pellets were eaten, the mouse was returned to an empty cage for 30 sec before the next trial was started. Every animal performed four trials per day. If the mouse turned into the baited arm first, the trial was counted as successful. After 7 days of training, the first probe trial was performed. During the probe trial, the training start arm was blocked, and mice entered the maze at the end of the initially blocked arm. If the mouse turned in the arm that was correct during training, the mouse was considered a “place learner”. If the mouse turned in the direction that was not correct during training, the mouse was considered a “response learner”. After another 7 days of training, the second probe trial was performed. In total every mouse performed 4x habituation, 56x training, and 2x probe trials.

### Automated home-cage discrimination and reversal learning test (CognitionWall)

An automated home-cage platform (Phenotyper, Noldus) was used to monitor spontaneous behavior as well as discrimination and reversal learning in absence of human intervention. Mice were monitored by video and tracked throughout the complete experiment using Ethovision 16 (Noldus). Up to 16 mice were tested in parallel. Before the start of the experiment, animals were single-housed in regular cages on white cellulose ALPHA-DRI bedding for 4-5 days and habituated to the rewarded sugar pellets (Dustless Precision Rodent Pellets, F05684, Bio-Serv, Flemington, NJ) with regular feed and water *ad libitum*. The lights were controlled by the automated home cage, the dark cycle was from 6 pm to 6 am. On the day of the experiment start, mice were transferred to the Phenotyper cages (L=30 x W = 30 x H = 35 cm) and habituated to the novel environment for 6 hours. The CognitionWall was introduced about 30 min (~4:00 pm) before the start of the discrimination learning experiment (~4.30 pm). The Cognition Wall, an opaque wall with three holes, allowed for the mice to pass through to retrieve food pellets, as previously described (Remmelink et al. (2016)). Water was provided *ad libitum* throughout the protocol. Standard feed was absent from the cage during the trial, but a “free” pellet reward was dispensed to create the association between Cognition Wall entry and pellet reward. During a 48-hour discrimination learning (DL) test, mice were trained to discriminate the left entrance hole as the “correct” hole and received a pellet reward. “Correct” hole entries were detected by the software and automatically triggered pellet dispensation. DL success was measured by rate of establishing a preference for the rewarded entrance. If the mouse did not reach the criterion of >100 dispensed pellets within 48 h, it was withdrawn from the study and returned to its home cage. After the DL phase, the reversal learning (RL) phase was conducted within the following 48 hours. During RL, the rewarded entrance shifted from “left” to “right” entrance. The rate of a shift in preference for the new entrance was used as a measure for reversal learning. During DL and RL, a pellet reward was dispensed for every fifth entry through the correct hole (FR5 schedule of reinforcement) to avoid the accumulation of non-consumed pellet rewards in the cage Remmelink et al. (2016). Mice did not have to make five consecutive correct entries. The results of the Cognition Wall experiments were independently analyzed by Sylics (Synaptologics BV).

### Acquisition, extinction, renewal of cued and contextual fear learning

Mice were tested for fear learning and memory in fear conditioning boxes (MED Associates). The experiment had these three phases: acquisition on day 1, fear extinction on day 2-4, and a test for context-dependent/renewal learning on day 5.

During the training on day 1, each mouse was put in the test chamber designated as context A (standard grid floor with isopropanol/simple green scent) within a sound-attenuating box and allowed to explore for 3 min. Then, the mice were exposed to a 75 dB 2800 Hz pure tone for 30 s that co-terminated with a 2 s scrambled foot shock (0.5 mA). Mice received 2 additional shock-tone pairings, with an 80 s pause between each pairing.

During the fear extinction phase on day 2-4, mice were placed into a modified chamber arranged as context B (black A-frame insert, white floor, ethanol/windex scent) for a test of extinction of the cued fear response in absence of the foot shock. After 3 minutes in the chamber, the animal was presented with a massed extinction protocol, where the same auditory tone from the acquisition day (30 s 75 dB 2800 Hz) was presented 20 times separated by 5 s. During the context-dependent/renewal learning phase on day 5, the mouse was returned to the original acquisition context conditioning chamber arranged as context A. After the exploration phase, the 75 dB 2800 Hz pure tone was presented 3x for 30 s per repetition using the protocol used for acquisition with the shock omitted. Freezing, defined as complete immobility except that necessitated by breathing was scored using the Video Freeze software (activity threshold 19 for 1 s). During the extinction phase tone presentations were binned by 5 tone presentations to facilitate graphing and analysis.

### Statistics

The experimenters were blinded for the mouse genotype during behavioral testing. Statview (SAS, Cary, NC, RRID:SCR_017411), Microsoft Excel (Redmond WA, USA, RRID:SCR_016137), Igor Pro 8.04 (Wavemetrics, Lake Oswego, OR, USA, RRID:SCR_000325), Prism 9 (GraphPad, San Diego, CA, USA, RRID:SCR_002798), and R (version 3.6.3) were used for data analysis. For the behavioral data, two-way or repeated measures analysis of variance (ANOVA) were used to determine effects of genotype and sex, followed by separate analyses for males and females, to determine genotype effects within each sex. Post-hoc comparisons were conducted using Fisher’s Protected Least Significant Difference (PLSD) tests only when a significant F value was found in the ANOVA. Within-genotype comparisons were used to determine side preference in the 3-chamber test and quadrant selectivity in the water maze. If no significant F value was found in the ANOVA for sex, animals were pooled by genotype.

For the T maze, no side bias was detected using graphical inspection. Success rate to enter goal arms, was analyzed by a Repeated Measures Proportional Odds Logistic Regression model using the repolr package (version 3.4, https://CRAN.R-project.org/package=repolr). Daily success rate from day 1 to day 14 was fit using generalized estimating equations with sex, genotype, day, and their interactions as well as an AR (1) covariance structure to reflect temporal correlation. The T maze probe trial results as well as strategy transitions from probe trial 1 to probe trial 2 were analyzed using a Fisher’s exact test. For all comparisons, significance was set at p<0.05.

## Results

### Loss of GABA co-transmission from ACh neurons does not affect general health, anxiety-like behavior, or motor and sensory skills in mice

The co-transmission of ACh and GABA has been described previously in the hippocampus, striatum, mPFC, lateral septum, as well as the retina (Lee et al. (2010), Saunders et al. (2015a), Saunders et al. (2015b), Sethuramanujam et al. (2016), Takacs et al. (2018), Lozovaya et al. (2018), Obermayer et al. (2019), Granger et al. (2020), Hunt et al. (2022)). Although these reports provided many insights into how ACh/GABA co-transmission is embedded into the individual circuits or contributes to pathogenicity in disease, the role of GABA co-transmission with ACh as it relates to circuit function remains unknown. Therefore, the primary goal of our experiments was to assess how a systemic loss of GABA co-transmission from ACh neurons affected the behavioral performance of mice during a battery of standardized behavioral tests (Table 1). We found that in absence of GABA co-transmission from ACh neurons, mice showed no impairments in general health, motor activity (locomotion in a simple environment, rearing, swimming, motor coordination), sensory abilities (vision, hearing, olfaction), anxiety-like behavior, sensorimotor gating, fear memory acquisition, or fear extinction (Fig. 1-1 to 4-1, Table 1-1). However, we found several genotype or sex-dependent effects on more complex cognitive processes, such as: social novelty preference, spatial memory, context-dependent locomotion, competing learning strategies, and fear renewal (Fig. 1-5, Table 2-1 to 6-1).

**Figure 1:**
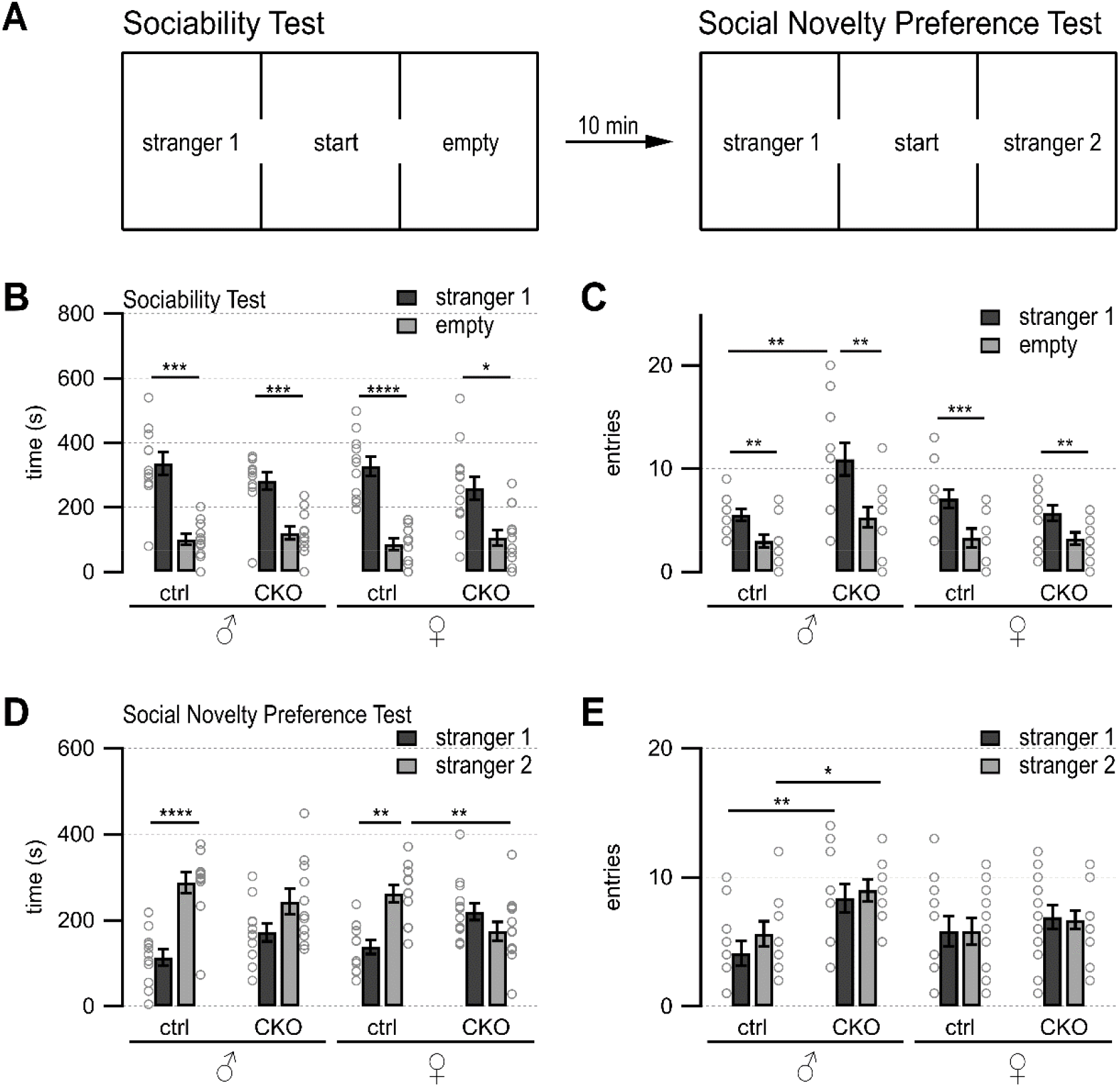
3-Chamber Test. **Loss of GABA co-transmission leads to a loss of social novelty preference in mice as well as increased chamber entries in males but does not affect sociability.** A, Schematic showing setup of sociability and social novelty preference test. Mice habituated for 10 min in the cage with both side chambers empty. Before the sociability test, an unfamiliar sex-matched mouse (stranger 1) was introduced into the left chamber. After 10 min of the sociability test, before start of the social novelty preference test, an unfamiliar sex-matched mouse (stranger 2) was introduced into the right chamber. Time spent in outer chambers and chamber entries were measured. B-C, Sociability test results. Time spent in (B) and entries (C) into side chambers shown for male, female control (ctrl) and CKO mice. D-E, Social novelty test results. Time spent in (D) and entries (E) into side chambers shown for male, female control (ctrl) and CKO mice. Values represent mean ± SEM. Individual data points are depicted as open circles. Repeated measures ANOVA followed by Fisher’s protected least-significant difference tests (*p<0.05, **p<0.01, ***p<0.001, ****p<0.0001). See Table 2-1 for data.

### GABA co-transmission from ACh neurons is required for social novelty preference independent of sex and exploratory behavior in males

Many psychiatric diseases, such as autism, bipolar disorder, and schizophrenia, are accompanied by impaired social abilities (Pietropaolo et al. (2011), Sidhu et al. (2014), Moy et al. (2004), Moy et al. (2009), Moy et al. (2013), Win-Shwe et al. (2021), O’Tuathaigh et al. (2007), Olaya et al. (2018), Nakazawa et al. (2019), Brunner et al. (2015), Carter et al. (2011), Jaramillo et al. (2017), Kim et al. (2017), Lu et al. (2018)). Using the three-chamber choice test (Fig. 1 A), we assessed whether a loss of GABA co-transmission from ACh neurons leads to impairments in the natural inclination of mice to investigate a novel social stimulus. Sociability, or the preference for an animal to investigate another animal compared with an empty chamber, was unaffected by loss of GABA co-transmission from ACh neurons (Fig. 1 B, main effect of side, males, F(1,20)=54.92, p<0.0001; females, F(1,22)=33.5, p<0.0001).

During the social novelty preference phase, only the ctrl but not the CKO mice showed significant social novelty preference as assessed by investigation times (Fig. 1 D, genotype x side, males F(1,20)=20.46, p=0.0002; females F(1,22)=11.23, p=0.0029). However, male CKO mice transitioned about twice as often into the two outer chambers during the sociability phase (Fig. 1 C, genotype, F(1,20)=9.46, p=0.006; genotype x side, F(1,20)=4.58, p=0.0448). Chamber entries during the social novelty preference phase were increased in male CKO mice, as well (Fig. 2 E, genotype, F(1,20)=11.44, p=0.003). Taken together, these data indicate that loss of GABA co-transmission from ACh neurons impaired social novelty preference, but not sociability, independent of sex. Additionally, we saw increased chamber entries in male CKO mice.

**Figure 2:**
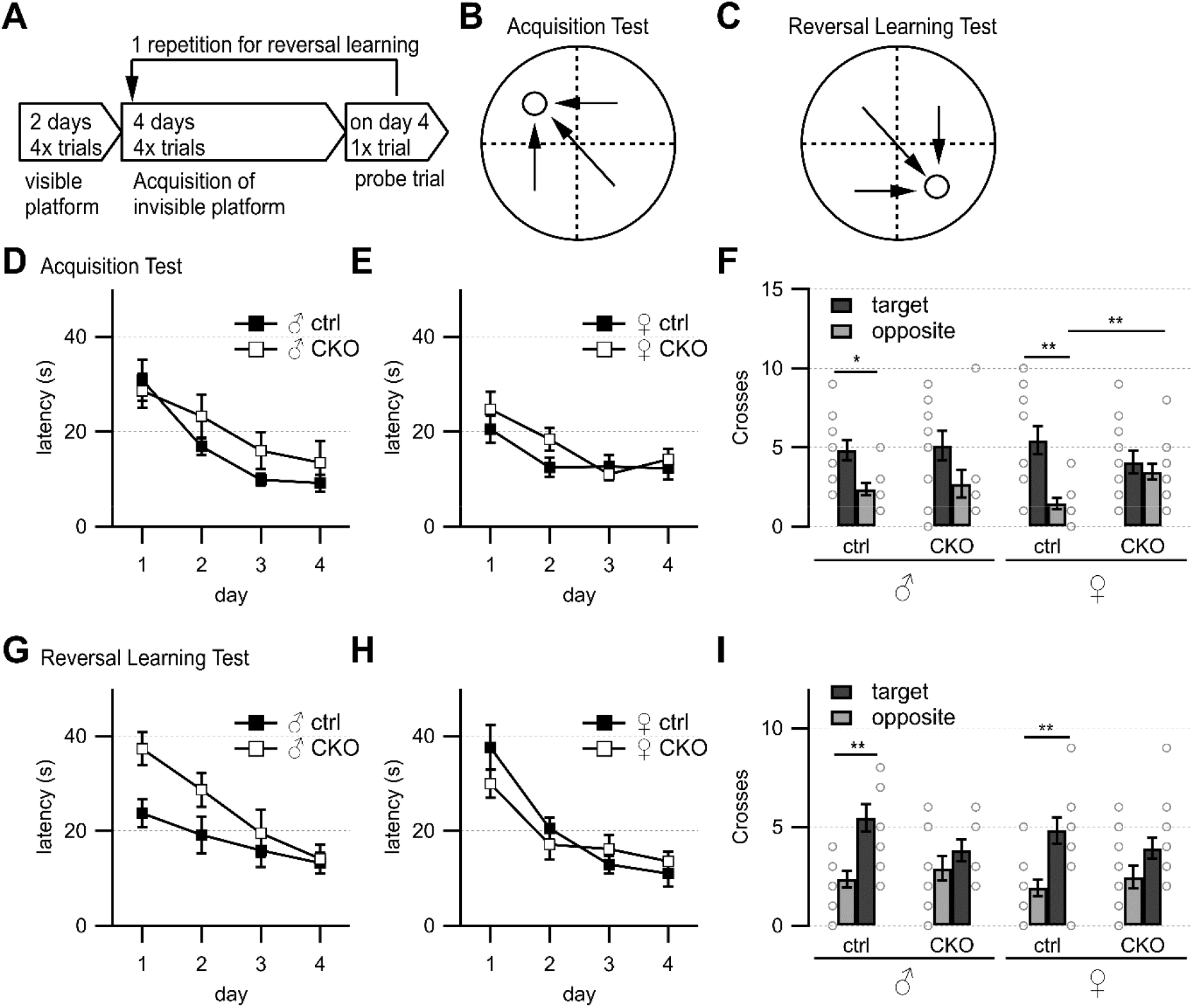
Morris water maze. **Loss of GABA co-transmission leads to spatial memory impairments.** A, Timeline of Morris water maze experiment. Visible platform test, acquisition test using hidden platform followed by 1 probe trial without platform, reversal learning test with hidden platform in opposite quadrant and 1 probe trial without platform. B, Water maze setup for acquisition test. Visible/hidden platform placed at circle. Start points chosen in arbitrarily random order. Theoretical escape pathway to hidden platform indicated by arrows. C, Water maze setup for reversal test with hidden platform placed in opposite quadrant. D-E, Latency to reach hidden platform during acquisition learning for males (D) and females (E). F, Probe trial results after acquisition learning phase in absence of platform. Counts of target and opposite quadrant crosses. G-H, Latency to reach hidden platform during reversal learning for males (D) and females (E). I, Probe trial results after reversal learning phase in absence of platform. Counts of swimming path crosses over platform area in target quadrant and corresponding area in opposite quadrant. Values represent mean ± SEM. Individual data points are depicted as open circles. Repeated measures ANOVA followed by Fisher’s protected least-significant difference (*p<0.05, **p<0.01). See Table 3-1 for data.

### GABA co-transmission from ACh neurons is required for spatial learning and memory

Previous work has shown that stimulation of septohippocampal ACh neuron terminals evoked biphasic ACh/GABA postsynaptic currents in oriens lacunosum moleculare (OLM) interneurons, which play an important role in information flow through the hippocampus/entorhinal cortex circuitry (Takacs et al. (2018), Haam et al. (2018)). Thus, we wanted to test whether a loss of the co-transmitted GABA from ACh neurons affected hippocampal function. We used the Morris water maze to assess swimming ability, spatial learning and memory, as well as reversal learning (Fig. 2 A-C, Brandeis et al. (1989), Vorhees and Williams (2014), Hollup et al. (2001)). On day 1 of the visible platform test, female CKO mice needed significantly more time to escape (genotype, F(1,22)=9.46, p=0.0055; genotype x side, F(1,22)=7.33, p=0.0129, Table 3-1). However, on the following day, all groups learned the visible platform test similarly well with comparable swim speed (Tab. 3-1). The escape latencies and the number of training days were not significantly different for either sex or genotype during the hidden platform test (Fig. 2 D-E). During the first probe trial, ctrl mice showed a significant preference for swim path crossings over the platform area in the target quadrant compared to the corresponding area in the opposite quadrant (Fig. 2 F, genotype x quadrant, males, F(1,19)=6.72; p=0.0179, females, F(1,22)=10.53, p=0.0037). In contrast, CKO mice did not show a preference for the target quadrant over the opposite quadrant (Fig. 2 F). Furthermore, female CKO mice had significantly more crossings over the incorrect area in the opposite quadrant than female ctrl mice (Fig. 2 F, genotype x quadrant, F(1,22)=5.67, p=0.0264)

During the reversal learning phase, male CKO and female mice showed a tendency to escape more slowly onto the hidden platform (Fig. 2 G-H). However, all animals escaped to the hidden platform within the 15 s criterion at the end of the 4-day reversal learning phase. Although CKO mice failed to demonstrate preference for the target quadrant during the second probe (Fig. 2 I, males, main effect of quadrant, F(1,19)=16.9, p=0.0006; genotype x quadrant, F(1,19)=5.09, p=0.036; females, main effect of quadrant, F(1,22)=16.9, p=0.0005), the initial spatial memory deficit likely overshadowed any potential effects on reversal learning. Taken together, these data indicate that GABA co-transmission from ACh neurons is required for spatial memory.

### GABA co-transmission from ACh neurons is not required for reward learning but stabilizes usage of competing learning strategies

The results of the Morris water maze test were suggestive of spatial learning deficits in CKO mice while the locomotive hyperactivity in the 3-chamber test suggests possible striatal impairments involving inhibitory behavioral control. In order to assess whether these deficits affected learning, we utilized a classic T-maze task that can be solved via either a hippocampus-dependent place strategy or a striatum-dependent response strategy and utilizes probe tests to determine which strategy is dominant (Packard and McGaugh (1996)). Recently, impaired inhibition in the dentate gyrus has been implied in decreased spatial learning during foraging behaviors (Albrecht et al. (2022)). Because ACh release levels in the hippocampus and striatum can be used as predictors of which strategy is dominant, we hypothesized that the loss of GABA co-transmission may lead to changes in reward learning and learning strategy usage (Chang and Gold (2003)).

We did not find any differences in distance travelled during the two days of habituation (Table 4-1). During the 14 days of training, we found no sex differences, and no differences in success rate except for a slight deficit trend on day 9 after the first probe trial (Fig. 3 C, p=0.096, Repeated Measures Proportional Odds Logistic Regression with Bonferroni correction). The first probe trial revealed an overall preference for the response learning strategy but no significant differences between ctrl and CKO mice (Fig. 3 D, p=0.303, Fisher’s exact test). During the second probe trial, strategy preferences roughly followed those of the first probe trial, with no significant differences between ctrl and CKO mice (Fig. 3 E, p=0.294, Fisher’s exact test). We assessed the stability of the response vs. place strategy usage further by assigning every animal to one of these four groups (Fig. 3 F): place transitioner (“R➔P”), consistent place learner (“Place”), consistent response learner (“Response”), response transitioner (“P➔R”). There were significantly more consistent response learners among ctrl mice compared to CKO mice (p=0.0434, Fisher’s exact test). This indicates that ctrl mice quickly adopted a striatal learning strategy by probe trial 1 and persisted in this strategy until probe trial 2, whereas CKO mice were less likely to maintain a consistent strategy. Taken together, these data indicate that a loss of GABA co-transmission from ACh neurons caused only minor changes in reward-associated learning in the T maze. However, the loss of GABA co-transmission from ACh neurons caused an instability in hippocampal vs. striatal strategy selection.

**Figure 3:**
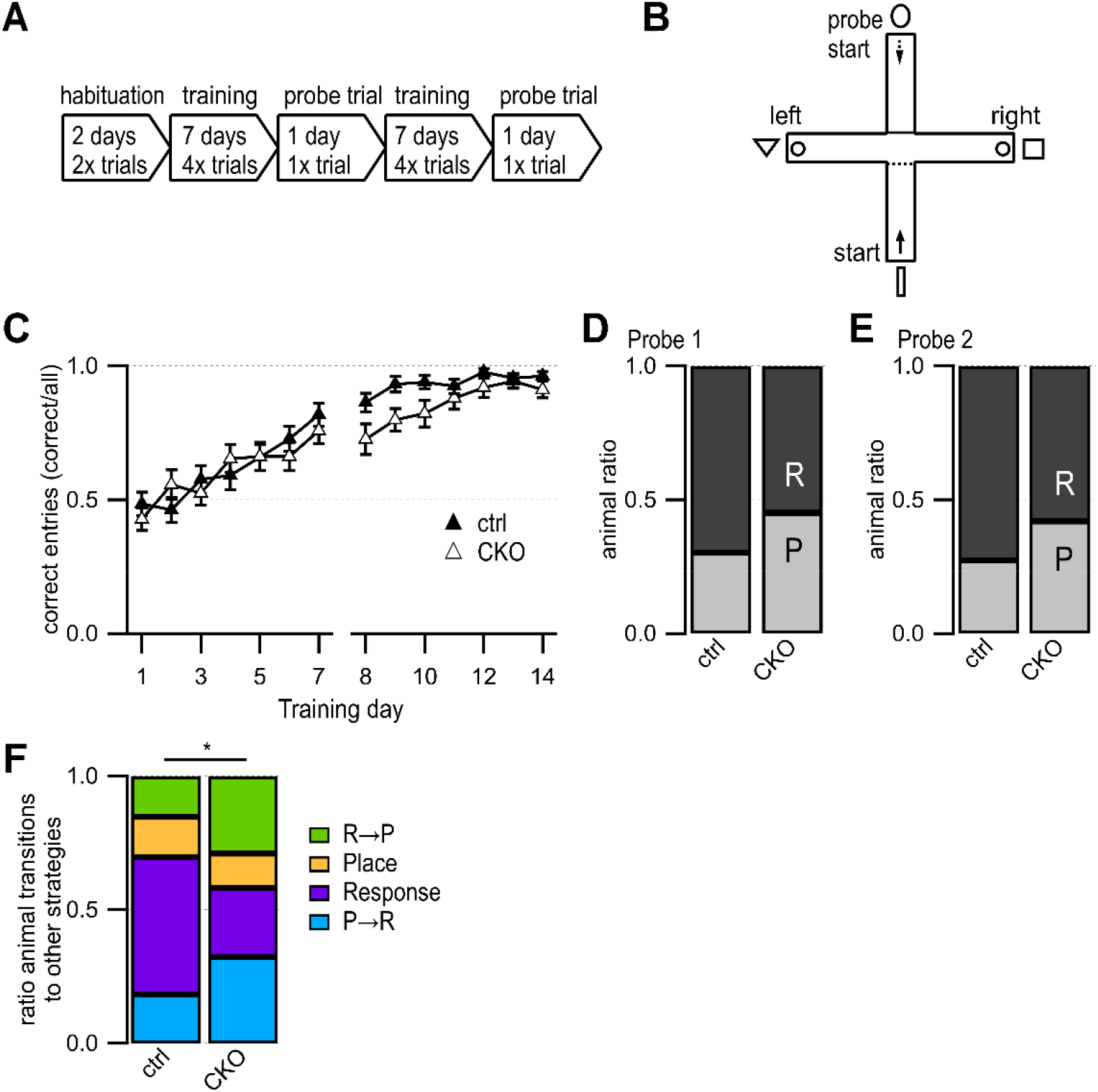
T Maze. **Loss of GABA co-transmission leads to changes in usage of competing learning strategies.** A, Timeline of T maze experiment. Habituation phase for 2 days, training phase for 7 days followed by 1 probe trial day. Repetition of training phase for another 7 days followed by 1 probe trial day. B, T maze setup. Animals start in south arm during habituation and training but from the north arm during probe trials. Visible fabric cues attached to the curtained walls outside of the maze. Food bowls (circle) were placed at ends of both goal arms. Start points for training indicated by continuous arrow and for probe trials by dashed arrow. C, Training success indicated as correct entries into goal arms for every training day. D-E, Results for Probe trial day 1 (D) and Probe trial day 2 (E) indicating the relative number of response (R) and place (P) learners. F, Animal ratios were sorted by learning strategy and strategy transitions from probe trial 1 to probe trial 2: place transitioner (R➔P, green), consistent place (orange), consistent response (purple), response transitioner (P➔R, turquoise). Values represent mean ± SEM. Repeated Measures Proportional Odds Logistic Regression model (C) and Fisher’s exact test (F) (*p<0.05). See Table 4-1 for data.

### GABA co-transmission from ACh neurons is not required for discrimination and reversal learning but regulates locomotion in males

We wanted to further assess spontaneous behaviors as well as learning performance, and cognitive flexibility of mice in the absence of human interference (Remmelink et al. (2016)). During a 4-day discrimination and reversal learning paradigm in an automated home-cage (Fig. 4 A), mice were trained to discriminate between rewarded or unrewarded response options followed by a reversal learning phase. During the discrimination learning (DL) phase, mice had to enter the correct (left) entrance into the CognitionWall to receive food rewards (Fig. 4 B). After the first two days, the reversal learning (RL) phase started with the correct response option changed from the left to the right entrance.

**Figure 4:**
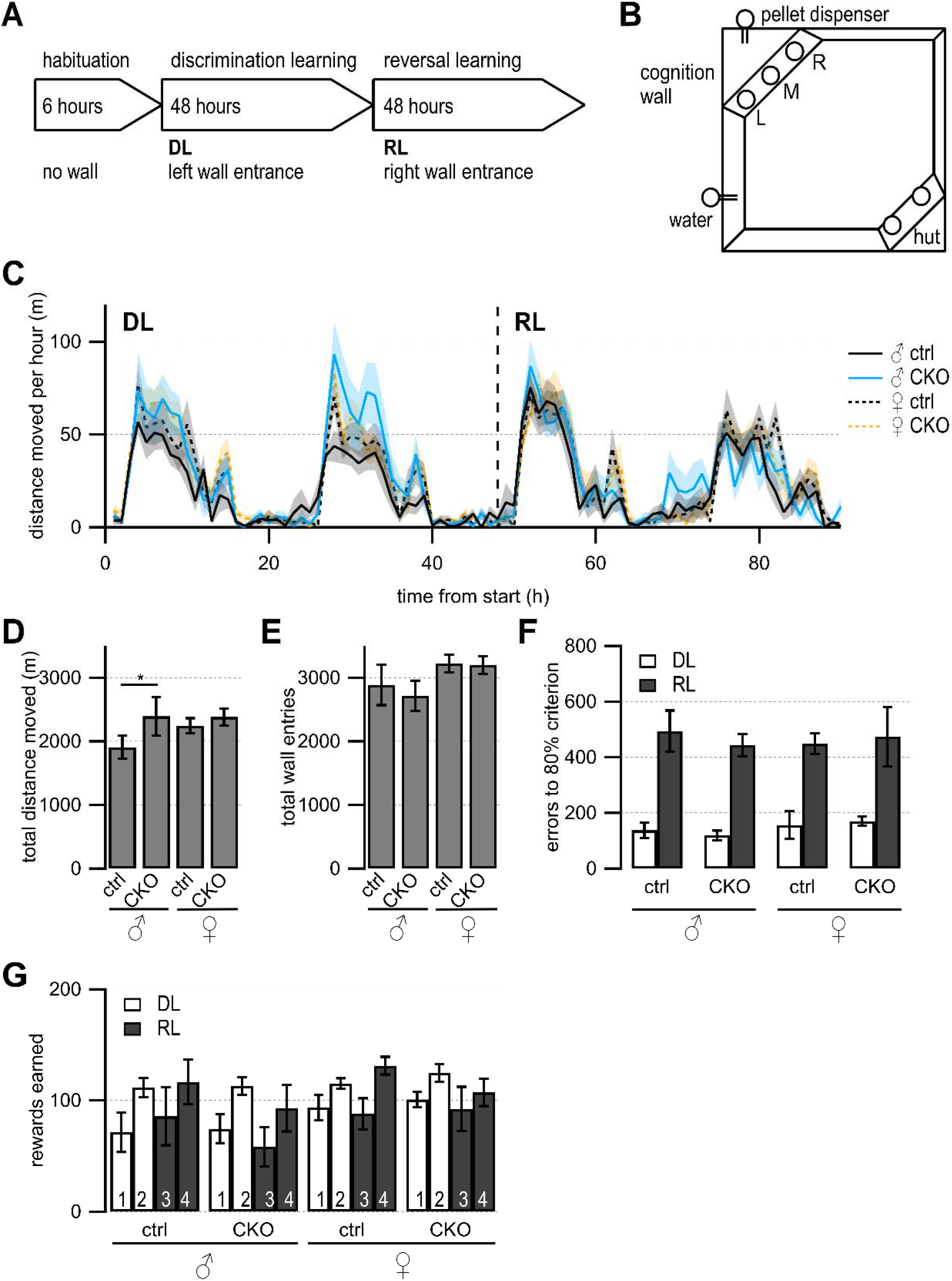
CognitionWall automated cage discrimination and reversal learning test. **Loss of GABA co-transmission from ACh neurons results in increased locomotion in males but no impairments in discrimination or reversal learning.** A, Timeline of CognitionWall experiment. Habituation phase for 6 hours without CognitionWall. Introduction of CognitionWall ~30 min before start of discrimination learning phase (DL) for 2 days followed by reversal learning phase (RL) for another 2 days. During DL, animal trained to enter left wall entrance for food reward. During RL, animal trained to enter right wall entrance for food reward. B, CognitionWall setup. Hut for shelter in bottom right, water bottle in bottom left, CognitionWall in top left covering the food dispenser. Left (L), middle (M), right (R) entrance of CognitionWall. C, Total distance moved during experiment per 1 hour bin. D, Total distance moved during experiment. E, Total wall entries during experiment. F, Numbers of error entries before reaching 80% learning criterion during DL and RL. G, Number of reward pellets earned per day during DL and RL. Values represent mean ± SEM. Two-way ANOVA followed by Fisher’s protected least-significant difference (*p<0.05). See Table 5-1 for data.

First, we looked at circadian rhythmicity of activity by assessing the distance moved over time (Fig. 5 C). All mice showed comparable activity patterns with the highest movement activity during the dark cycle. We investigated animal activity by comparing total distance moved and found that male CKO mice moved significantly more than male ctrl mice during the experiment (Fig. 4 D, males, F(89, 1260)=27.66, p=0.0409). Female mice showed a higher activity compared to male ctrl mice (Fig. 4 D, females, F (3, 3060)=113.5, p=0.0106). However, the total wall entry count was not significantly different between any of the groups (Fig. 4 E). During DL and RL, all groups reached the 80% learning criterion within a similar number of errors (Fig. 4 F) comparable to C57BL/6J mice, (Remmelink et al. (2016)). We didn’t find differences in rewards earned per day (Fig. 4 G).

**Figure 5:**
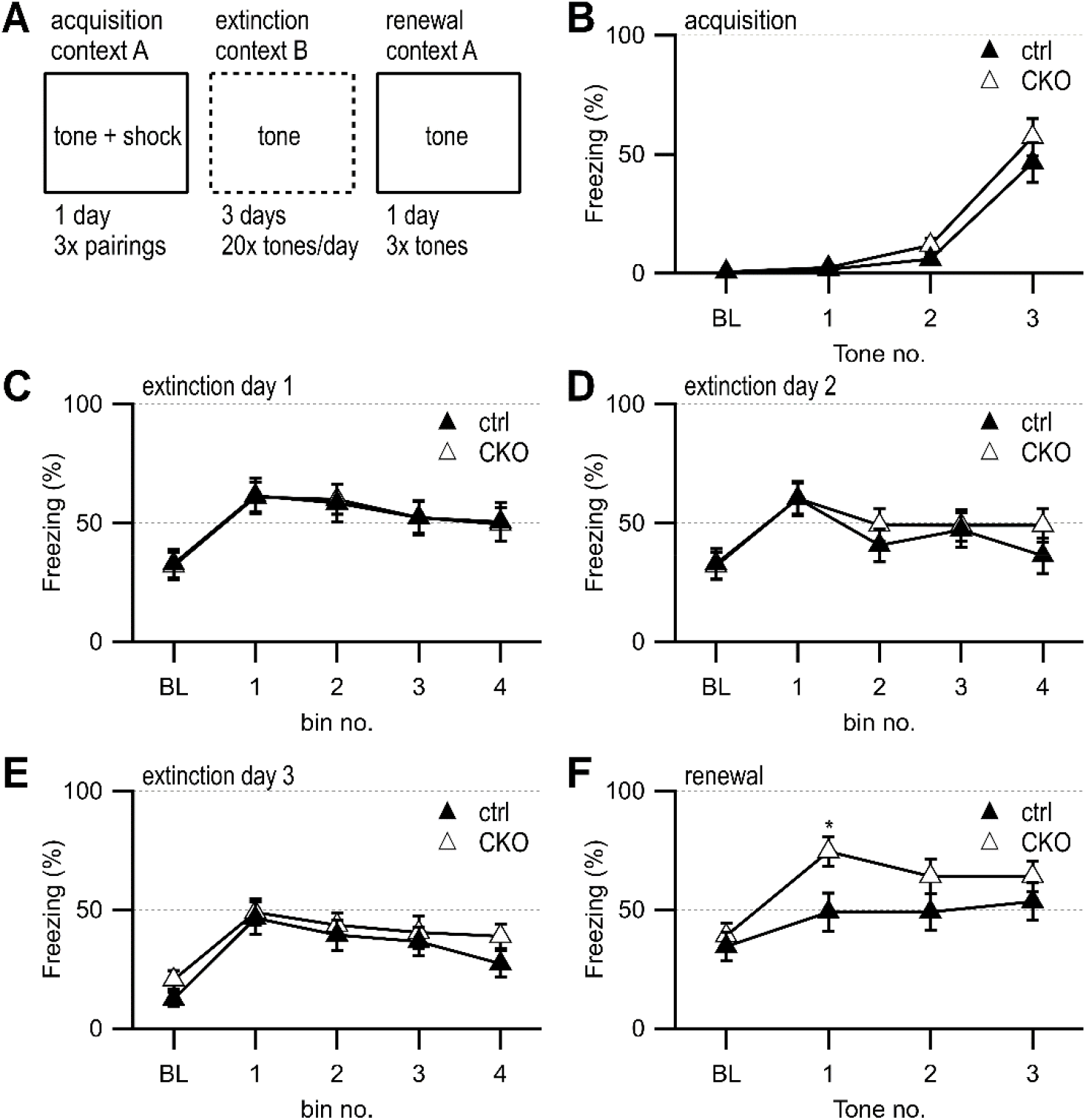
Fear Extinction. **Loss of GABA co-transmission from ACh neurons leads to transient increased freezing during contextual fear renewal.** A, Timeline of fear conditioning experiment. Acquisition of fear memory on day 1 by 3x pairing of 75 dB 2800 Hz pure tone with 2 s 0.5 mV foot shock in context A. During 3 days of extinction, animal was presented with 20 tones/day in context B. During the renewal of the fear memory, animal was presented with 3 tones in context A. B, Percent of animals freezing during acquisition phase at baseline (BL) and per tone/foot shock pairing. C-E, Percent of animal freezing during extinction phase at baseline (BL) and per 5 tone bin on day 1 (C), day 2 (D), and day 3 (E). F, Percent of animals freezing during extinction phase at baseline (BL) and per tone presentation. Values represent mean ± SEM. Two-way ANOVA followed by Fisher’s protected least-significant difference test (*p<0.05). See Table 6-1 for data.

Taken together, these data indicate that the loss of GABA co-transmission from ACh neurons does not affect discrimination or reversal learning in an automated home-cage experiment. However, we detected higher locomotive activity in male CKO mice without affecting wall entry or rewards earned. We therefore propose GABA co-transmission from ACh neurons may be required for locomotive or exploratory behaviors in males.

### GABA co-transmission from ACh neurons is not required for acquisition or extinction of fear memories, but for context-dependent renewal of fear to a discrete auditory cue

During the initial behavioral assessment, CKO mice did not exhibit substantial alterations in acquisition and retention of fear memories (Fig. 4-1). The other behavioral deficits in CKO mice, however, pointed towards disturbed communication between the hippocampus, the mPFC, and the striatum after the loss of GABA co-transmission from ACh neurons. Given the prominent role the mPFC plays in fear learning and prior evidence for GABA/ACh co-transmission in mPFC circuitry function, particularly counteracting roles of infralimbic cortex (IL) and prelimbic cortex (PL), we wanted to further assess whether fear extinction might be affected (Likhtik and Johansen (2019), Marek et al. (2018), Vasquez et al. (2019), Saunders et al. (2015a), Obermayer et al. (2019), Granger et al. (2020)).

We found no sex differences and again observed normal acquisition (Fig. 5 B) as well as a similar reduction in freezing during repeated tone presentations during fear extinction (Fig. 5 C-E). However, when placed back into the original training context for a test of context-dependent fear renewal, the CKO mice showed a transient elevation of freezing during the first tone presentation (Fig. 5 F, a priori planned comparison of tone 1, genotype: F(1,35) = 5.293, p = 0.027). These findings indicate that GABA co-transmission from ACh neurons may contribute to fear renewal.

## Discussion

Here, we studied the role of GABA co-transmission from ACh neurons in behavior. In the absence of GABA co-transmission from ACh neurons, mice showed substantial impairments in social and spatial memory, less stability in learning strategy usage, as well as context-dependent fear renewal. Furthermore, we discovered a task-specific increase in locomotion exclusively in males.

### Impairments in social novelty preference

The 3-chamber test revealed a loss in social novelty preference in CKO mice, but not a loss in sociability. The hippocampal regions CA2 – and CA1 to some extent - process novelty and social information (Alexander et al. (2016), Okuyama et al. (2016)). The CA2 region receives inputs from the supramammillary nucleus, which is activated by novelty and influences θ oscillations during exploration of novel stimuli (Ito et al. (2009), Vertes et al. (2004), Jeewajee et al. (2008)). In addition, social novelty preference requires ACh stimulation from basal forebrain neurons to the hippocampus where α7 nAChRs on GABAergic neurons lead to disinhibition in CA2 (Pimpinella et al. (2021), Nacer et al. (2021)). Other signaling cascades are involved in CA2 function and social memory, such as glucocorticoids, enkephalins, and neuropeptides like oxytocin and vasopressin (Pagani et al. (2015), Raam et al. (2017), McCann et al. (2021), Leroy et al. (2022)). Hippocampal sharp-wave ripples and output to the mPFC through γ oscillations depend on CA2 activity (Joo and Frank (2018), Alexander et al. (2018)). Although it is unclear whether GABA co-transmission is directly required for γ oscillations, blocking GABA transmission was sufficient to block sharp-wave ripples *in vitro* (Takacs et al. (2018)).

Proper mPFC function and hippocampus/mPFC crosstalk are required for social behaviors and disturbed in mouse models for study of autism, Alzheimer’s disease, or other brain dysfunctions (Riedel et al. (2009), Liu et al. (2020), Huang et al. (2020), Bicks et al. (2020), Franklin et al. (2017), Phillips et al. (2019), Gordon (2011), Johnson et al. (2013), Kim et al. (2017), Liang et al. (2018), Li et al. (2015), Poppe et al. (2019)). Given the ACh/NMDA receptor interplay in mPFC glutamatergic signaling and social behaviors, GABA co-transmission may contribute to neuron tuning and disruption in autism models in mice and higher-order mammals (Cools and Arnsten (2022), Avale et al. (2011), Okada et al. (2021), Sun et al. (2022), Liang et al. (2018) Okuyama et al. (2016), Brigman et al. (2009), Finlay et al. (2015)). Taken together, the impairment of social novelty preference in CKO mice may be caused by impaired signaling in the hippocampus, the mPFC, or other brain regions.

### Impairments in spatial learning and memory

The Morris water maze revealed spatial memory impairments in CKO mice. The hippocampus/entorhinal cortex circuitry is essential for allocentric navigation which is required for spatial learning and memory (O’Keefe (1978), Leutgeb et al. (2005), Buzsaki and Moser (2013), Chersi and Burgess (2015)). Hippocampal function requires well-timed ACh modulation (Hasselmo and McGaughy (2004), Gu and Yakel (2011)). During exploration, hippocampus and associated brain regions exhibit large type 1 θ oscillations involving M1 mAChR on hippocampal pyramidal neurons as well as α7 nAChR on OLM interneurons (Gu et al. (2017), Gu et al. (2020)). Because OLM interneurons receive ACh/GABA co-transmission from MS/DBB inputs, co-transmitted GABA may also modulate hippocampal spike timing and output through OLM interneurons (Takacs et al. (2018), Haam et al. (2018)).

The formation and consolidation of spatial memories requires additional hippocampal and mPFC interplay through VIP^+^ neurons (Maviel et al. (2004), Lee et al. (2019), Malik et al. (2022)). Although only ~10% of mPFC VIP^+^ neurons are ACh^+^, GABA co-transmission may modulate disinhibition or cause other effects in the mPFC (Obermayer et al. (2019), Granger et al. (2020)). Taken together, loss of GABA co-transmission from ACh neurons likely impaired spatial memory through altering hippocampus, mPFC, or both.

### Task-specific increases in locomotive activity in males

We observed increased locomotion in male CKO mice only in the 3-chamber test and the automated home-cage assay. Consistent with the literature, however, female ctrl mice were generally more active than male ctrl mice in the automated home-cage assay (Caldarone et al. (2008), Rosenfeld (2017), Holcomb et al. (2022), Warncke et al. (2021), Stevanovic et al. (2022)). Changes in locomotion have been reported in drug abuse models, in response to disturbed striatal DA transport and receptor function, and in patients with bipolar disorders or schizophrenia potentially through ventral striatum nAChR signaling (Moy et al. (2013), Singer et al. (2012), Perry et al. (2009), Moreno et al. (2013), DeLong and Wichmann (2015), Amitai et al. (2014), Jerlhag et al. (2006)).

In the striatum, ACh^+^ neurons relay signals between striosome and matrix neurons (Brimblecombe and Cragg (2017), Crittenden et al. (2017), Inoue et al. (2016)). Striatal ACh^+^/GABA^+^ neurons receive strong cortical inputs, are involved in the pause response and sensitive to inhibition while ACh neurons are less sensitive (Canales and Graybiel (2000), Miyamoto et al. (2018), Davis et al. (2018), Lozovaya et al. (2018)). Therefore, the loss of GABA co-transmission from ACh neurons may affect locomotor activity by disrupting striatal integration of mPFC signals, balance of striatal compartments, or feedback to the mPFC (Saunders et al. (2015b)).

### Role of GABA co-transmission in competing learning strategies

The animal’s ability to employ place or response learning to solve a goal-directed task has intrigued researchers for a long time (Goodman (2020), Chersi and Burgess (2015)). Overall, mice showed a preference for striatal response learning in the T maze, but CKO mice were less consistent in their strategy choices.

During striatal learning, the dorsolateral striatum undergoes changes in neuron firing, cell signaling, as well as epigenetic modifications (Aosaki et al. (1994b), Zhang and Cragg (2017), Jog et al. (1999), Kheirbek et al. (2009), Malvaez et al. (2018)). These changes likely enable the synchronization between striatum and hippocampus leading to increased mPFC activity (Doeller and Burgess (2008), Doeller et al. (2008), Goldenberg et al. (2020)). Given the importance of ACh neurons in striatal learning and function, loss of GABA co-transmission may disrupt corticostriatal inhibition, striatal output, strengthen striatal extinction mechanisms, or reduce synchrony between the hippocampus, the mPFC, and striatum (Aosaki et al. (1994a), Chang and Gold (2003), Lozovaya et al. (2018), Goldenberg et al. (2020), Fleming et al. (2022)). We, therefore, conclude that GABA co-transmission is a minor regulator of response learning with possible relevance to altered learning strategies in addiction, autism spectrum disorders, schizophrenia, or PD (Graybiel and Rauch (2000), Redgrave et al. (2010)).

### Role of GABA co-transmission in fear renewal

Disturbances in the underlying neural circuitry of fear response are associated with phobias or post-traumatic stress disorder (PTSD) (Herry et al. (2010)). We did not see changes in acquisition or fear extinction but found transient increases in context-dependent fear renewal in CKO mice.

Disturbances in ACh signaling in neurons or glia in hippocampus, amygdala and mPFC were found in cue encoding, spatial processing, aversive learning, extinction, and fear renewal (Likhtik and Johansen (2019), Yanpallewar et al. (2022), Zhang et al. (2021), Lotfipour et al. (2013), Lotfipour et al. (2017), Kutlu et al. (2018), Titus et al. (2019), Mineur et al. (2020), Chen et al. (2020), Mooney-Leber et al. (2021), Kellis et al. (2020), Yanpallewar et al. (2022), Miguelez Fernandez et al. (2021), Oliveros-Matus et al. (2020)). For example, feed-forward inhibition from the ventral hippocampus to PL is sufficient to decrease contextual fear renewal, while feed-forward inhibition from ventral hippocampus to IL increases fear renewal (Marek et al. (2018), Vasquez et al. (2019)). Moreover, fear extinction and fear renewal are improved by GABA_B_R inhibition at the beginning of the extinction period independent of dorsal hippocampus or BLA (Adkins et al. (2021)). We, therefore, propose that GABA co-transmission from ACh neurons may inhibit fear renewal in mice by coordinating hippocampus-mPFC crosstalk. Given that the changes in fear responses were transient, GABA co-transmission from ACh neurons likely has only a minor role in regulating contextual fear renewal to a conditioned tone.

### Role of GABA Co-Transmission from ACh Neurons in the Brain

Taken together, our findings support the importance of GABA co-transmission from ACh neurons in the brain and agree with previously published work (see Fig. 6 for simplified model). Loss of GABA co-transmission from ACh neurons impaired hippocampal function likely due to alterations in OLM interneuron function (Takacs et al. (2018)). This may affect both memory formation and consolidation through dysregulation of the hippocampus-entorhinal cortex loop as well as synchronized oscillatory events associated with social or spatial information (Alexander et al. (2018), Gu et al. (2017), Gu et al. (2020), Haam et al. (2018)). Since oscillatory activities in different brain regions are highly correlated, asynchronous hippocampal output may negatively affect mPFC activity and output to downstream targets or feedback to the hippocampus (Lee et al. (2019), Malik et al. (2022)).

**Figure 6:**
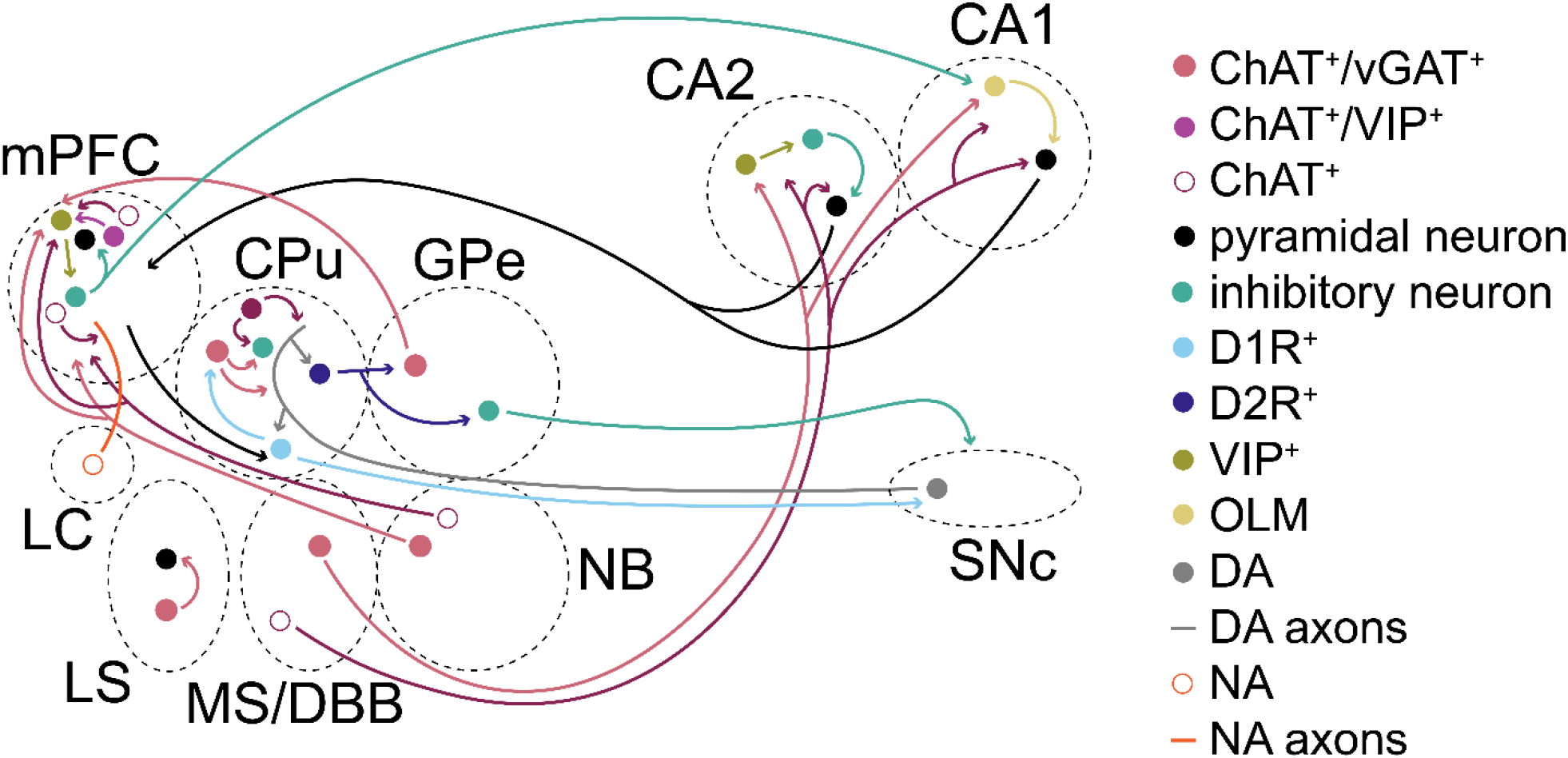
Summary/Simplified Model. **Simplified model of how ACh/GABA co-transmission is embedded in circuitry of cortical and subcortical structures.** ACh/GABA projections from the basal forebrain (MS/DBB and NB) form synapses in the hippocampus and mPFC, respectively. Basal ganglia structures such as CPu or GPe contain either local ACh/GABA interneurons or projection neurons projecting to the mPFC. The mPFC contains local ACh/GABA interneurons and receives noradrenergic (NA) inputs from the locus coeruleus (LC).

Striatal function was likely impaired by decreased cortical disinhibition through GABA^+^/ACh^+^ interneurons during the pause response of ACh neurons and may have implications in PD or Tourette syndrome (Lozovaya et al. (2018), Lennington et al. (2016)). The role of GABA co-transmission from ACh neurons in GPe and NB in circuit function remains unclear (Saunders et al. (2015b), Saunders et al. (2015a)). While the GPe output to the mPFC could provide negative feedback from the striatum through the indirect pathway during habit formation, NB may add spike timing refinement in mPFC L1. Similarly, local mPFC GABA co-transmission mostly targets L1 interneurons (Obermayer et al. (2019), Granger et al. (2020)). Given the sparsity, one may wonder about the physiological relevance in adult mice. However, GABA co-transmission may further shape the circuit during critical windows in development, as recently shown for GABAergic transmission in the mPFC or motor cortex (Bicks et al. (2020), Steinecke et al. (2022)). Higher-order mammals, such as rats, could potentially co-transmit more GABA from ACh neurons (Bayraktar et al. (1997), Obermayer et al. (2019), Dienel et al. (2021)).

In the future, it may be beneficial to assess how GABA co-transmitting ACh neurons are integrated into the brain circuitry. By studying anatomy of cell distributions and synaptic projections as well as the functional consequences for pre- or postsynaptic targets, GABA co-transmission from ACh neurons may be better understood. The behavioral relevance of GABA co-transmission could be further elucidated by restricting its loss to specific brain regions or cell populations. Lastly, future studies may help to determine the developmental role of GABA co-transmission during neurocircuit formation.

## Author Contributions

conceptualization: R.O.G., J.L.Y, J.D.C., S.S.M.

data curation: R.O.G., K.M.H., B.J.B., S.A.F.

analysis: R.O.G., S.S.M., J.D.C., P.W.L. figures: R.O.G.

writing: R.O.G.

## Funding

This research was supported by the Intramural Research Program of the NIH (NIH grant number Z01ES090089 to J.L.Y.), National Institute of Environmental Health Sciences, and by the Center on Compulsive Behaviors, NIH via NIH Director’s Challenge Award (R.O.G.), and the UNC Intellectual and Developmental Disabilities Research Center (NICHD; P50 HD103573; PI: Joseph Piven).

## Acknowledgments

We thank for statistical support by Dr. Min Shi at NIEHS and Dr. Sandra McBride, Dr. Guanhua Xie, and Dr. Kathryn Konrad (contract #: GS-00F-173CA / 75N96022F00055). We are grateful for assistance with animal husbandry/feeding/care provided by Scotty Dowdy, Maria Barrientos, Steven Butler, Rodriguez Sutton, as well as the NIEHS Comparative Medicine Branch. We thank Makayla Wood for technical assistance. We thank Drs. Shiyi Wang, Serena M. Dudek, as well as Yakel lab members for comments on the manuscript.

## Conflict of interest

The authors declare that the research was conducted in the absence of any commercial or financial relationships that could be construed as a potential conflict of interest.

## Notes

### Competing Interest Statement

The authors have declared no competing interest.

