## Supplemental_Material for "Loss of GABA co-transmission from cholinergic neurons impairs behaviors related to hippocampal, striatal, and medial prefrontal cortex functions"

**Supplemental Data**

**Supplemental Methods**

**Elevated plus maze**

Mice performed one 5-min trial on a plus maze with two walled arms (the closed arms, 20 cm in height) and two open arms. The maze was elevated 50 cm from the floor, and the arms were 30 cm long. Every mouse was placed in the maze center (8 cm x 8 cm) and allowed to freely explore the maze. Measures were taken of time on, and number of entries into, the open and closed arms.

**Open field test**

To assess exploratory activity in a novel environment, mouse movement was tracked during a one-hour trial in an open field chamber (41 cm x 41 cm x 30 cm) crossed by a grid of photobeams (VersaMax system, AccuScan Instruments). During the trial, counts were taken of the number of photobeams broken in 5-min intervals, measuring both locomotor activity (total distance traveled) and vertical rearing movements. An index of anxiety-like behavior was determined by the time spent in the center region.

**Rotarod test**

Mice were assessed for motor coordination on an accelerating rotarod (Ugo Basile, Stoelting Co., Wood Dale, IL). On the first day, mice were given three trials, with 45 seconds between each trial. After 48 hours, another three trials were given to evaluate consolidation of motor learning. Each trial was started at an initial value of 3 revolutions per minute (rpm), with a progressive increase to a maximum of 30 rpm across the maximum trial length of five minutes. The latency to fall from the top of the rotating barrel was assessed during the trials.

**Marble-bury assay**

The mice were examined in a Plexiglas cage located in a sound-attenuating chamber with ceiling light and fan. The cage was filled with 5 cm of corncob bedding, and 20 black glass marbles (14 mm diameter) arranged in an equidistant 5 X 4 grid on top of the bedding. Subjects were given access to the marbles for 30 min. After that time, the number of buried marbles was counted (2/3 of the marble covered by bedding).

**Acoustic startle test**

The mice were assessed regarding auditory function, reactivity to environmental stimuli, and sensorimotor gating. These functions are required for the reflexive startle response following a sudden, loud sound. Test results were startle magnitude and pre-pulse inhibition, which occurs when a weak prestimulus leads to a reduced startle in response to a subsequent louder noise. Mice were transferred into individual Plexiglas cylinders within larger, sound-dampening chambers (San Diego Instruments SR-Lab system). A piezoelectric transducer was located underneath every chamber to record the force of the startle responses. Every chamber also contained a ceiling light, fan, and a loudspeaker for the acoustic stimulation. A digital sound level meter (San Diego Instruments) was used to measure background sound levels (70 dB) and calibrate the acoustic stimuli. After a five-min habituation period, animals underwent a total of 42 trials from 7 different trial types: no-stimulus trials, trials with the acoustic startle stimulus (40 msec; 120 dB) alone, and trials in which a pre-pulse stimulus (20 ms; either 74, 78, 82, 86, or 90 dB) occurred 100 msec before the onset of the startle stimulus. The startle amplitude was assessed for each trial during a 65-ms sampling window. For the analysis, each subject's data for levels of pre-pulse inhibition (PPI) at each pre-pulse sound level was determined as:

$$PPI=100-\frac{response amplitude (pre­pulse stimulus and startle stimulus)}{response amplitude (startle stimulus only)}*100$$

**Buried food test**

Palatability of an unfamiliar food (Froot Loops, Kellogg Co., Battle Creek, MI) was observed and consumption was documented for every mouse before the olfactory test. All food was removed from the home cage about a day before the test. During the test, each mouse was placed in a large, clean tub cage (46 cm L x 23.5 cm W x 20 cm H), filled with paper chip bedding (3 cm deep), and allowed five minutes exploration time. After the exploration phase, the animal was taken out of the cage, and one Froot Loop was buried in the cage bedding. During the test phase, the animal was returned to the cage and had 15 minutes to find the buried food. Latency to find the food reward was obtained to compare the olfactory abilities of the animals.

**Supplemental Material**

Figure 1-1 – Weight, elevated plus maze, Rotarod

Figure 2-1 – Open field test

Figure 3-1 – Pre-pulse inhibition

Figure 4-1 – Fear conditioning

Table 1-1 – Marble burying assay and buried food test and other assays (refers to Fig. 1-1 to 4-1)

Table 2-1 – 3-chamber test (refers to Fig. 1)

Table 3-1 –Morris water maze (refers to Fig. 2)

Table 4-1 – T maze (refers to Fig. 3)

Table 5-1 – CognitionWall (refers to Fig. 4)

Table 6-1 – Fear conditioning (refers to Fig. 5)

**Figure 1-1 – Weight, elevated plus maze, Rotarod**

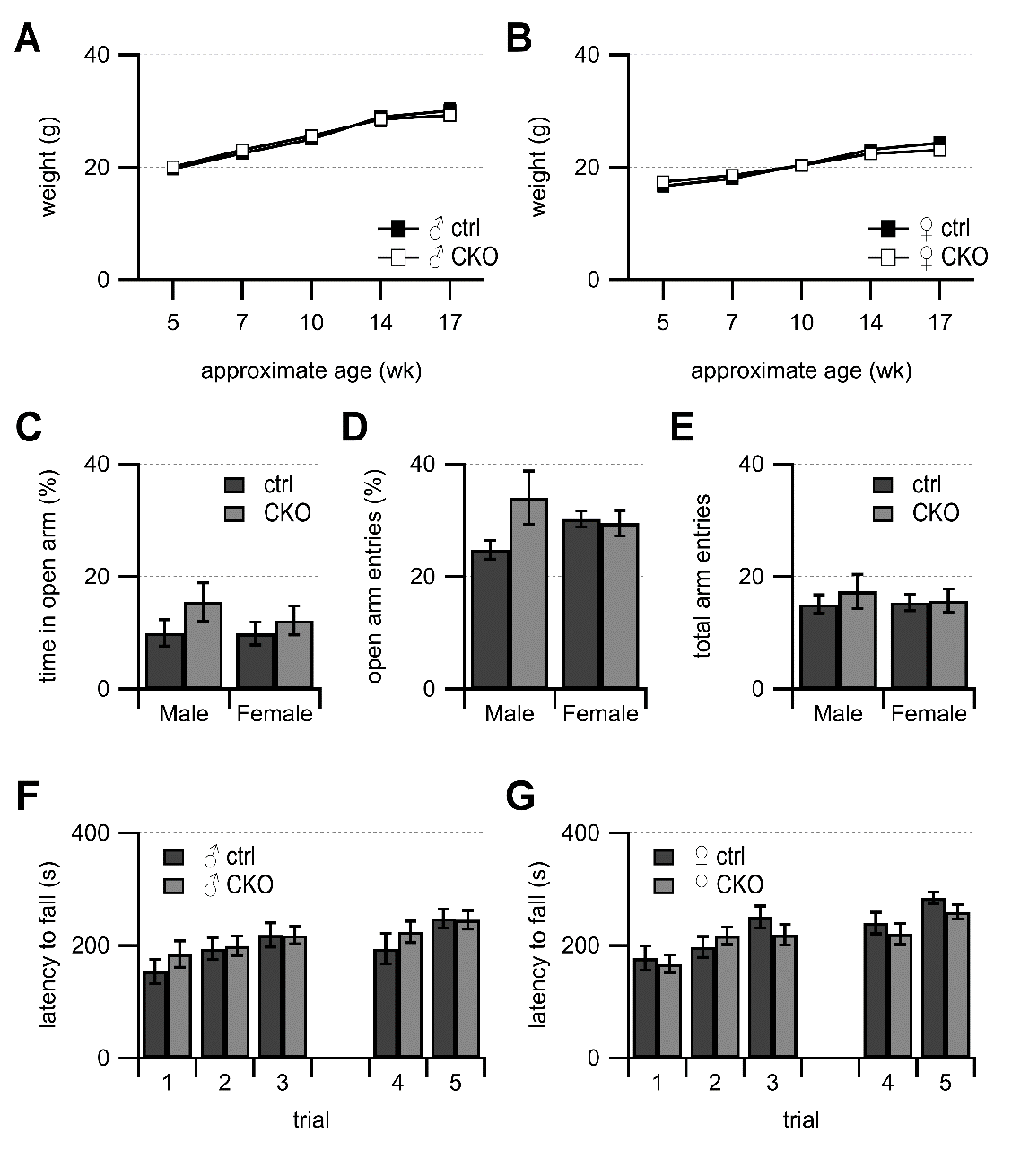

**Figure 1-1: Loss of GABA co-transmission from ACh neurons does not affect body weight, anxiety-like behavior, or motor coordination.** A-B, Animal weights for male (A) and female (B) mice during behavioral phenotyping. C-E, Relative time spent in open arm (C), relative open arm entries (D), and total arm entries (E) during 5 min elevated plus maze. F-G, Evaluation of motor coordination on an accelerating rotarod of male (F) and female mice (G). Maximum trial length 300 s. Trials 4 and 5 were given 48 hours after the first 3 trials. Values represent means ± SEM.

**Figure 2-1 – Open field test**

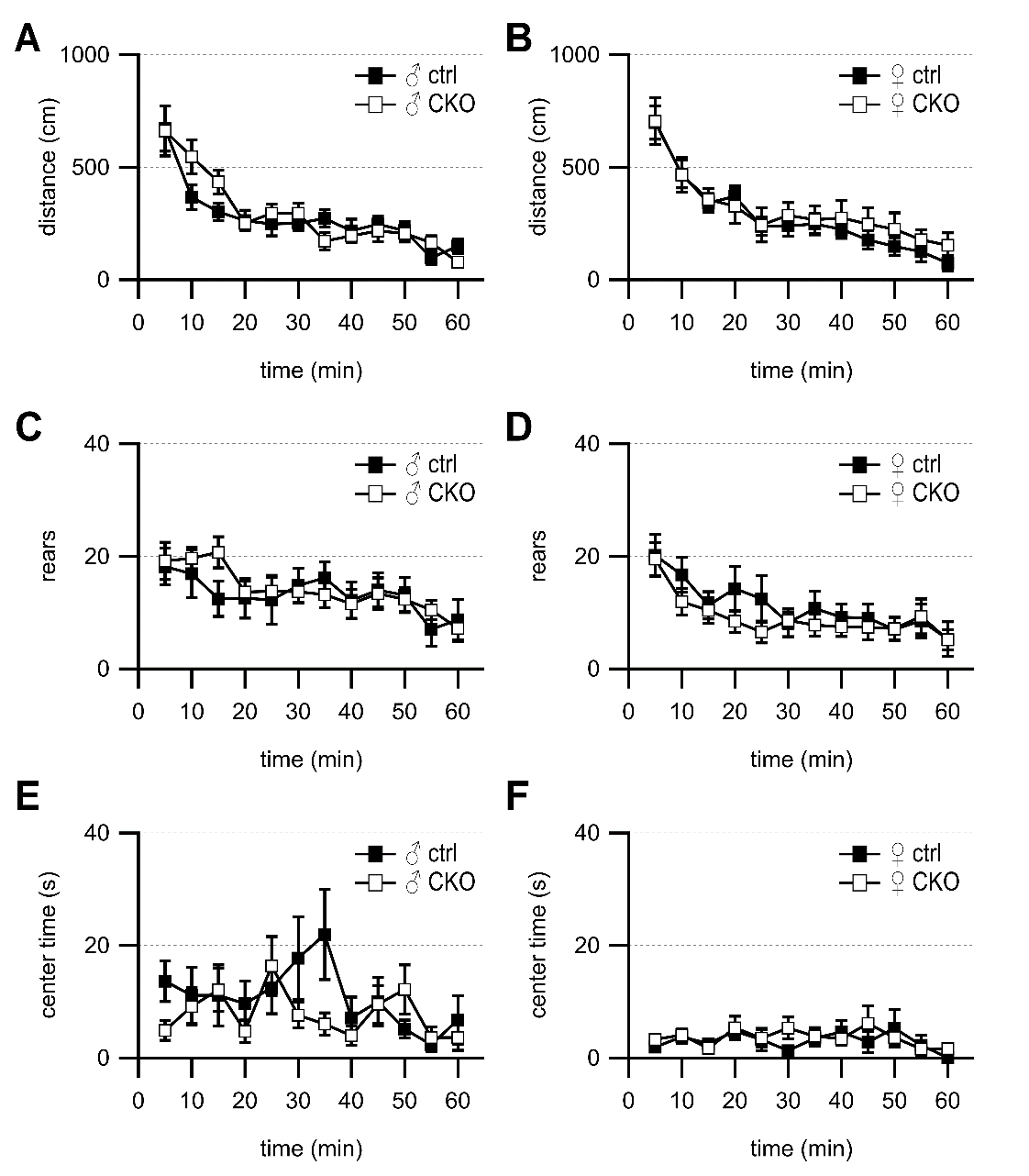

**Figure 2-1. Loss of GABA co-transmission from ACh neurons does not affect locomotor activity, rearing movements, or time spent in center region during an open field test.** A-B, Distance moved per 5-min bin for males (A), and females (B). C-D, Number of rears per 5-min bin for males (C) and females (D). E-F, Time spent in the center of chamber for males (E) and females (F). Values represent mean ± SEM.

**Figure 3-1 – Pre-pulse inhibition**

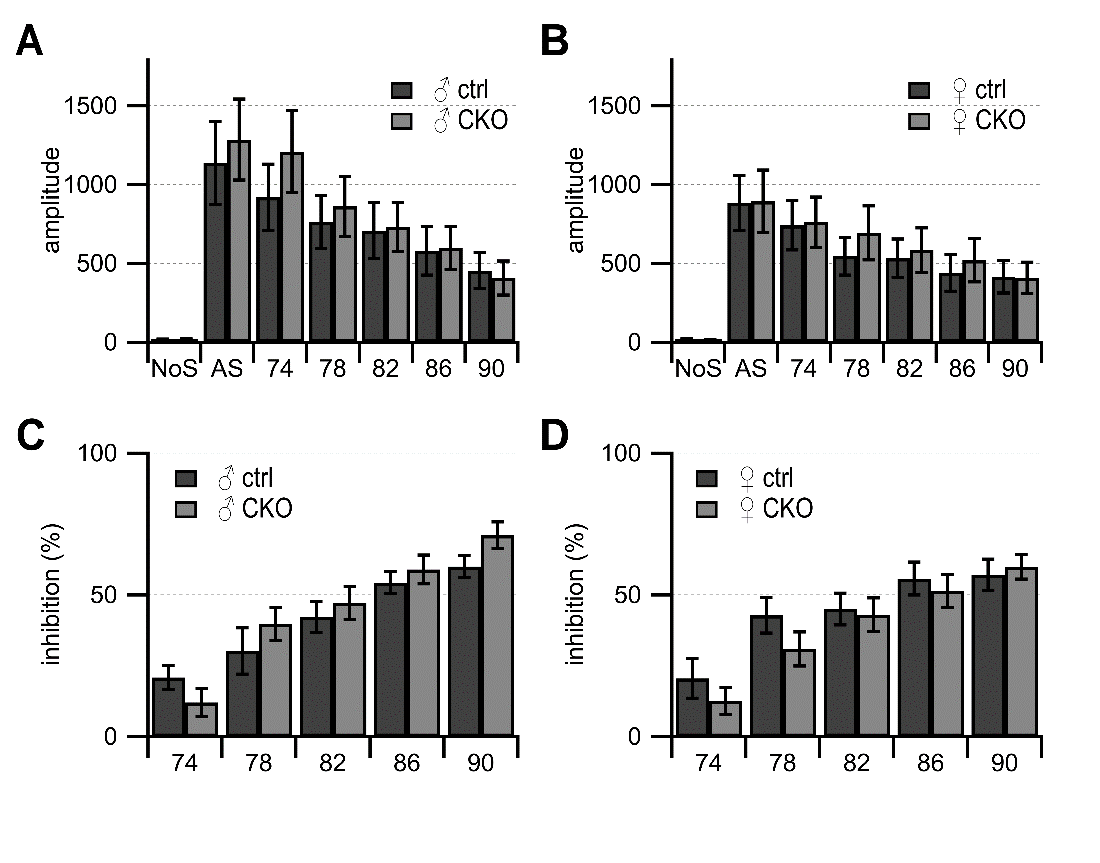

**Figure 3-1.**  **Loss of GABA co-transmission from ACh neurons does not affect magnitude of startle responses or pre-pulse inhibition.** A-B, Startle amplitude for males (A), and females (B). No auditory stimulus (NoS) trials, acoustic startle stimulus alone (AS; 120 dB), and pre-pulses at 74-90 dB. C-D, Pre-pulse inhibition for males (C) and females (D). Values represent mean ± SEM.

**Figure 4-1 – Fear conditioning**
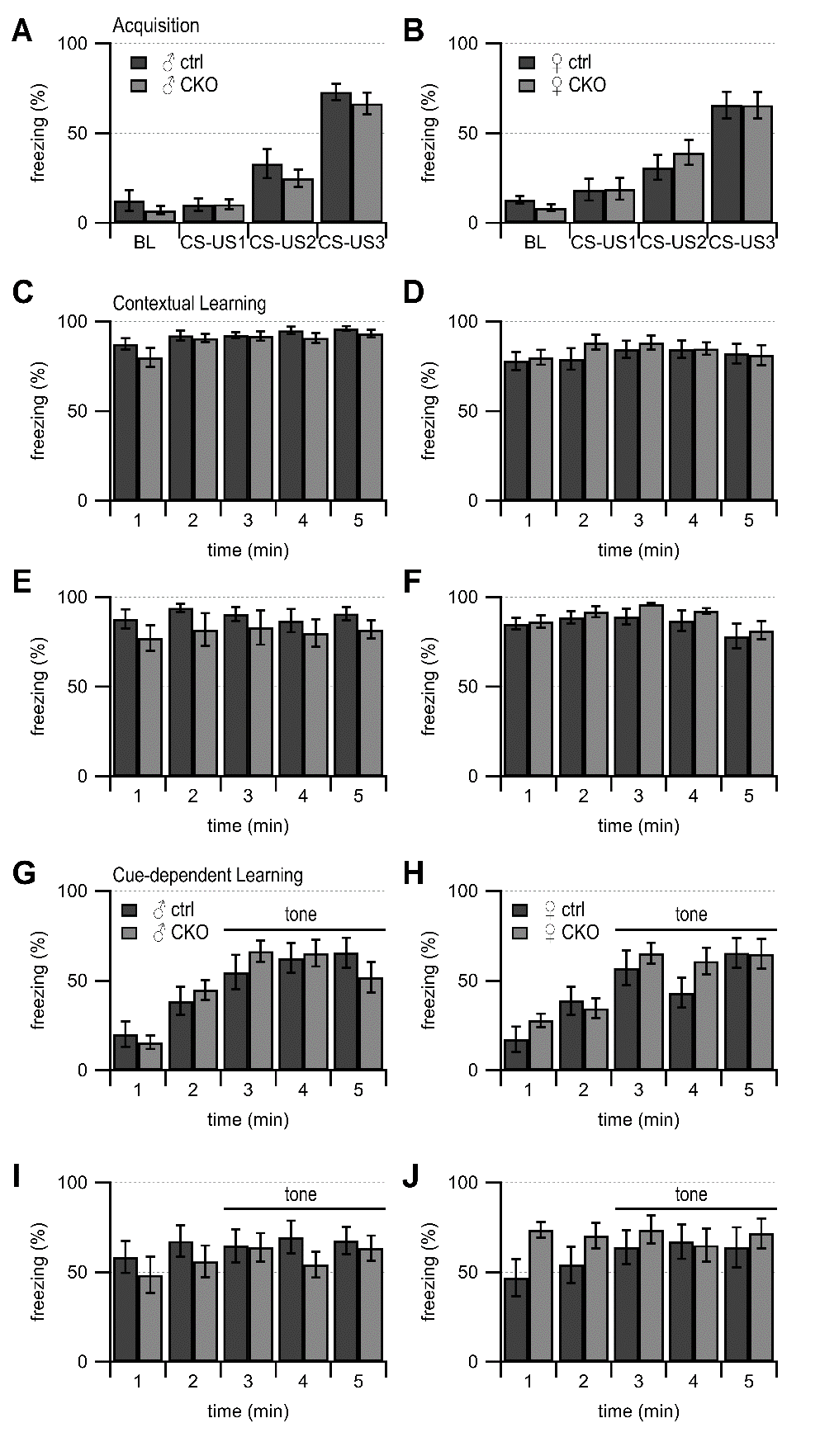

**Figure 4-1. Loss of GABA co-transmission from ACh neurons does not affect acquisition of conditioned fear memories, contextual, or cue-dependent learning.** A-B, Acquisition of fear memory for males (A) and females (B). Baseline freezing during a 2-min period before the presentation of tone-shock pairings (BL). Conditioned stimulus (CS, 30 s, 80 dB tone); Unconditioned stimulus (US, 2 s 0.4 mA foot shock) presented in the final 2 s of the tone. 3x tone-shock pairings during the training session, with 80 s between each pairing. C-F, The first 5-min contextual learning test conducted in the fear conditioned chambers 24 hr after the training session on Day 1 for males (C) and females (D). Contextual learning test 2 was conducted 2 weeks following Test 1 (E-F). G-J, The first cue-dependent learning test conducted 24 hr after the first contextual learning test for males (G) and females (H). Conditioned stimulus was presented 2 min after mice were placed in the modified conditioned fear chambers. Cue-dependent learning test 2 was conducted 2 weeks following Test 1 (I-J).

**Table 1-1 – Marble burying assay and buried food test and other assays (refers to Fig. 1-1 to 4-1)**

|  | **Males** | | | | **Females** | | | |
| --- | --- | --- | --- | --- | --- | --- | --- | --- |
|  | **ctrl** | | **CKO** | | **ctrl** | | **CKO** | |
|  | **mean** | **SEM** | **mean** | **SEM** | **mean** | **SEM** | **mean** | **SEM** |
| **Weight** | | | | | | | | |
| 5 weeks | 19.7 | 0.9 | 20.0 | 0.6 | 16.7 | 0.9 | 17.4 | 0.3 |
| 7 weeks | 22.5 | 0.6 | 23.0 | 0.6 | 18.0 | 0.6 | 18.5 | 0.4 |
| 10 weeks | 25 | 0.7 | 25.6 | 1.0 | 20.4 | 0.6 | 20.3 | 0.3 |
| 14 weeks | 28.9 | 1.0 | 28.5 | 1.1 | 23.1 | 0.5 | 22.4 | 0.4 |
| 17 weeks | 30.0 | 1.2 | 29.2 | 0.9 | 24.3 | 0.6 | 23.0 | 0.4 |
| N= | 11 |  | 11 |  | 11 |  | 13 |  |
| **Olfactory test** | | | | | | | | |
| Latency to find buried food (s) | 40 | 15 | 73 | 26 | 98 | 31 | 60 | 19 |
| % finding food | 100 |  | 82 |  | 91 |  | 92 |  |
| N= | 11 |  | 9 |  | 10 |  | 12 |  |
| **Marble-burying assay** | | | | | | | | |
| # buried in 30 min | 18 | 0.3 | 17 | 0.5 | 18 | 0.6 | 17 | 0.6 |
| N= | 11 |  | 11 |  | 11 |  | 13 |  |
| **Elevated plus maze** | | | | | | | | |
| Time open arm (s) | 34.6 | 6.1 | 42.5 | 9.3 | 28.7 | 6.4 | 33.6 | 7.0 |
| Open arm entries (%) | 12.4 | 2.3 | 15.4 | 3.4 | 9.9 | 2.0 | 12.2 | 2.6 |
| Total arm entries | 15.1 | 1.7 | 17.4 | 3.0 | 15.4 | 1.4 | 15.7 | 2.1 |
| N= | 11 |  | 11 |  | 11 |  | 13 |  |
| **Rotarod** | | | | | | | | |
| Latency to fall trial 1 (s) | 153.7 | 21.6 | 184.8 | 23.3 | 177.4 | 21.6 | 166.8 | 16 |
| Latency to fall trial 2 (s) | 194.2 | 19.3 | 199.3 | 17.5 | 197.4 | 18. 5 | 217.4 | 15.6 |
| Latency to fall trial 3 (s) | 218.5 | 21.7 | 218.3 | 15.3 | 250.5 | 19.6 | 218.8 | 18.2 |
| Latency to fall trial 4 (s) | 194. 3 | 27.3 | 224.1 | 19.1 | 239.7 | 19.3 | 220.3 | 18.8 |
| Latency to fall trial 5 (s) | 248.1 | 16.5 | 245.6 | 16.7 | 284. 4 | 10.5 | 259.7 | 12.7 |
| N= | 11 |  | 11 |  | 11 |  | 13 |  |
| **Open field test** | | | | | | | | |
| Distance (cm) 5 min | 672.0 | 100.7 | 661.0 | 110.8 | 698.8 | 74.7 | 704.8 | 102.8 |
| Distance (cm) 10 min | 366.4 | 54.1 | 546.8 | 75.7 | 469.5 | 61.7 | 466.1 | 76.7 |
| Distance (cm) 15 min | 302.4 | 38.3 | 435.4 | 52.9 | 340.9 | 41.3 | 356.5 | 48.7 |
| Distance (cm) 20 min | 262.5 | 46.3 | 250.6 | 25.9 | 370.5 | 45.6 | 326.6 | 77.1 |
| Distance (cm) 25 min | 247.7 | 52.3 | 295.3 | 39.7 | 237.7 | 40.4 | 243.6 | 76.4 |
| Distance (cm) 30 min | 251.6 | 33.0 | 295.1 | 44.3 | 239.9 | 45.9 | 286.3 | 56.8 |
| Distance (cm) 35 min | 272.3 | 38.2 | 171.5 | 39.4 | 247.5 | 47.4 | 266.5 | 61.5 |
| Distance (cm) 40 min | 217.5 | 51.4 | 195.0 | 29.6 | 224.7 | 40.8 | 274.1 | 77.5 |
| Distance (cm) 45 min | 247.0 | 35.0 | 216.9 | 46.6 | 176.9 | 40.2 | 247.2 | 72.0 |
| Distance (cm) 50 min | 218.5 | 41.0 | 206.3 | 38.4 | 148.5 | 38.9 | 224.3 | 73.0 |
| Distance (cm) 55 min | 97.5 | 29.1 | 164.2 | 31.0 | 124.3 | 43.9 | 177.8 | 45.2 |
| Distance (cm) 60 min | 148.8 | 33.2 | 78.7 | 22.0 | 74.8 | 34.7 | 152.9 | 55.4 |
| Rears 5 min | 18.2 | 3.4 | 19.2 | 3.3 | 20.2 | 3.7 | 19.5 | 2.9 |
| Rears 10 min | 16.9 | 4.3 | 19.6 | 2.0 | 16.7 | 3.1 | 11.9 | 2.3 |
| Rears 15 min | 12.5 | 3.1 | 20.7 | 2.8 | 11.3 | 2.4 | 10.4 | 2.3 |
| Rears 20 min | 12.5 | 3.5 | 13.6 | 2.0 | 14.3 | 4.0 | 8.5 | 2.0 |
| Rears 25 min | 12.3 | 4.4 | 13.8 | 2.5 | 12.5 | 4.2 | 6.5 | 2.0 |
| Rears 30 min | 14.8 | 3.1 | 13.7 | 1.9 | 8.2 | 2.4 | 8.6 | 1.4 |
| Rears 35 min | 16.2 | 2.8 | 13.2 | 2.3 | 10.7 | 3.0 | 7.8 | 2.0 |
| Rears 40 min | 12.2 | 3.2 | 11.5 | 2.6 | 9.2 | 2.3 | 7.5 | 1.7 |
| Rears 45 min | 14.0 | 3.0 | 13.4 | 2.8 | 9.0 | 2.6 | 7.4 | 2.3 |
| Rears 50 min | 13.1 | 3.1 | 12.4 | 2.1 | 7.0 | 2.0 | 7.2 | 2.0 |
| Rears 55 min | 7.0 | 3.0 | 10.5 | 1.8 | 8.5 | 3.0 | 9.3 | 3.1 |
| Rears 60 min | 8.7 | 3.6 | 7.2 | 2.4 | 5.3 | 3.1 | 5.2 | 1.8 |
| Center (s) 5 min | 13.7 | 3.6 | 4.9 | 1.8 | 1.9 | 0.4 | 3.3 | 0.7 |
| Center (s) 10 min | 11.1 | 5.0 | 9.2 | 3.3 | 3.8 | 1.2 | 4.1 | 1.0 |
| Center (s) 15 min | 11.1 | 5.4 | 12.1 | 3.8 | 2.6 | 0.6 | 1.7 | 0.9 |
| Center (s) 20 min | 9.6 | 4.0 | 4.7 | 2.0 | 4.8 | 1.4 | 5.3 | 2.1 |
| Center (s) 25 min | 12.5 | 4.6 | 16.4 | 5.2 | 3. 3 | 2.0 | 3.5 | 1.6 |
| Center (s) 30 min | 17.7 | 7.3 | 7.6 | 2.3 | 1.3 | 0.6 | 5.3 | 1.9 |
| Center (s) 35 min | 22.0 | 8.0 | 6.0 | 2.0 | 3.5 | 1.5 | 3.9 | 1.6 |
| Center (s) 40 min | 7.1 | 3.7 | 4.0 | 1.7 | 4.6 | 2.1 | 3.5 | 1.2 |
| Center (s) 45 min | 10.0 | 4.3 | 9.5 | 3.3 | 2.8 | 1.8 | 6.1 | 3.2 |
| Center (s) 50 min | 5.2 | 1.6 | 12.2 | 4.4 | 5.3 | 3.4 | 3.6 | 1.4 |
| Center (s) 55 min | 2.3 | 1.3 | 3.6 | 1.9 | 2.2 | 1.8 | 1.6 | 1.2 |
| Center (s) 60 min | 6.8 | 4.3 | 3.6 | 2.3 | 0.1 | 0.04 | 1.7 | 0.7 |
| N= | 11 |  | 11 |  | 11 |  | 13 |  |
| **Pre-pulse inhibition** | | | | | | | | |
| Amplitude NoS | 17.4 | 1.9 | 18.3 | 1.8 | 17.1 | 3.0 | 14.1 | 1.2 |
| Amplitude AS | 1138.2 | 263.0 | 1286.0 | 256.4 | 882.2 | 174.7 | 893.8 | 198.8 |
| Amplitude 74 | 920.1 | 209.7 | 1210.2 | 258.9 | 741.7 | 155.5 | 761.8 | 160.0 |
| Amplitude 78 | 761.0 | 168.7 | 862.6 | 190.0 | 545.1 | 119.2 | 692.9 | 171.0 |
| Amplitude 82 | 707.0 | 177.4 | 732.4 | 154.6 | 532.4 | 122.2 | 584.9 | 141.2 |
| Amplitude 86 | 580.0 | 154.1 | 598.0 | 137.6 | 441.6 | 116.8 | 519.8 | 137.8 |
| Amplitude 90 | 453.9 | 113.8 | 407.1 | 107.7 | 414.3 | 102.3 | 408.5 | 100.2 |
| PPI 74 | 20.9 | 4.3 | 12.1 | 5.0 | 20.5 | 7.0 | 12.6 | 4.8 |
| PPI 78 | 30.2 | 8.2 | 39.8 | 5.9 | 42.8 | 6.2 | 30.9 | 6.0 |
| PPI 82 | 42.2 | 5.5 | 47.1 | 5.8 | 45.0 | 5.6 | 43.0 | 6.0 |
| PPI 86 | 54.3 | 3.9 | 59.0 | 5.1 | 55.8 | 5.8 | 51.5 | 5.8 |
| PPI 90 | 60.0 | 3.9 | 71.0 | 4.7 | 57.1 | 5.5 | 59.9 | 4.4 |
| N= | 11 |  | 11 |  | 11 |  | 13 |  |
| **Fear conditioning** | | | | | | | | |
| **Acquisition** | | | | | | | | |
| BL freezing (%) | 12.6 | 5.8 | 7.1 | 2.3 | 12.9 | 2.0 | 8.4 | 2.0 |
| Tone 1 (%) | 10.3 | 3.5 | 10.4 | 2.6 | 18.5 | 6.9 | 19.0 | 6.1 |
| Tone 2 (%) | 33.0 | 8.1 | 25.0 | 4.8 | 30.9 | 4.7 | 39.2 | 7.0 |
| Tone 3 (%) | 73.0 | 4.4 | 66.6 | 6.2 | 65.7 | 4.6 | 65.6 | 7.5 |
| **Contextual learning** | | | | | | | | |
| BL 1 freezing (%) | 87.5 | 3.1 | 80.1 | 5.3 | 78.0 | 5.0 | 80.0 | 4.1 |
| BL 2 (%) | 92.3 | 2.9 | 90.8 | 2.4 | 79.1 | 6.0 | 88.4 | 4.2 |
| T1 (%) | 92.6 | 1.6 | 91.8 | 2.7 | 84.5 | 4.8 | 88.2 | 4.0 |
| T2 (%) | 95.0 | 2.1 | 90.8 | 2.8 | 84.4 | 4.9 | 84.9 | 3.4 |
| T3 (%) | 96.1 | 1.3 | 93.3 | 2.3 | 82.1 | 5.4 | 81.2 | 5.6 |
| BL 1#2 (%) | 88.0 | 5.3 | 77.0 | 7.2 | 85.2 | 3.2 | 86.4 | 3.5 |
| BL 2#2 (%) | 94.0 | 2.2 | 81.9 | 9.1 | 88.7 | 3.3 | 91.9 | 3.1 |
| T1 #2 (%) | 90.5 | 3.8 | 83.0 | 9.5 | 89.0 | 4.4 | 96.0 | 0.9 |
| T2 #2 (%) | 86.8 | 6.5 | 80.1 | 7.6 | 87.0 | 5.7 | 92.3 | 1.6 |
| T3 #2 (%) | 90.8 | 3.8 | 82.0 | 5.0 | 78.1 | 6.9 | 81.5 | 5.2 |
| **Cue-dependent learning** | | | | | | | | |
| BL 1 freezing (%) | 20.2 | 7.105 | 15.572 | 3.7 | 17.2 | 4.1 | 27.9 | 8.0 |
| BL 2 (%) | 38.6 | 7.805 | 44.799 | 5.6 | 38.9 | 7.8 | 34.6 | 6.0 |
| T1 (%) | 54.7 | 9.605 | 66.399 | 5.8 | 57.1 | 8.5 | 65.3 | 7.0 |
| T2 (%) | 62.5 | 8.326 | 65.377 | 7.6 | 43.2 | 9.8 | 60.9 | 8.3 |
| T3 (%) | 65.6 | 8.372 | 51.867 | 8.4 | 65.5 | 7.5 | 64.9 | 5.2 |
| BL 1#2 (%) | 58.6 | 9.0 | 48.5 | 10.2 | 46.9 | 10.5 | 73.8 | 4.4 |
| BL 2#2 (%) | 67.5 | 8.6 | 56.0 | 8.9 | 54.2 | 10.1 | 70.4 | 7.0 |
| T1 #2 (%) | 64.8 | 9.3 | 63.9 | 8.0 | 64.0 | 9.4 | 73.8 | 7.9 |
| T2 #2 (%) | 69.7 | 9.1 | 54.3 | 7.2 | 67.1 | 9.6 | 65.1 | 9.1 |
| T3 #2 (%) | 67.8 | 7.7 | 63.4 | 7.1 | 63.9 | 11.3 | 71.7 | 8.4 |
| N= | 11 |  | 10 |  | 11 |  | 13 |  |

**Table 2-1 – 3-chamber test (refers to Fig. 1)**

|  | **Males** | | | | **Females** | | | |
| --- | --- | --- | --- | --- | --- | --- | --- | --- |
|  | **ctrl** | | **CKO** | | **ctrl** | | **CKO** | |
|  | **mean** | **SEM** | **mean** | **SEM** | **mean** | **SEM** | **mean** | **SEM** |
| **Habituation** | | | | | | | | |
| left side (s) | 156.9 | 25.9 | 116.2 | 20.0 | 149.8 | 14.1 | 150.3 | 29.5 |
| right side (s) | 154.5 | 28.9 | 153.8 | 19.5 | 160.5 | 21.2 | 143.0 | 21.6 |
| Entries left side | 7.8 | 1.4 | 8.0 | 2.1 | 9.0 | 1.4 | 7.2 | 0.9 |
| Entries right side | 7.2 | 1.1 | 10.4 | 1.6 | 8.6 | 1.4 | 7.2 | 1.1 |
| statistics^1^ | F(1,20)=3.74 | p=0.0676 |  |  |  |  |  |  |
| **Sociability** | | | | | | | | |
| stranger 1 (s) | 336.3 | 35.9 | 282.3 | 27.8 | 327.9 | 30.1 | 259.4 | 35.6 |
| empty (s) | 100.9 | 17.2 | 120.7 | 20.1 | 85.1 | 18.9 | 106.2 | 24.2 |
| statistics^2^ | F(1,20)=54.92 | p<0.0001 |  |  | F(1,22)=33.5 | p<0.0001 |  |  |
| Entries stranger 1 | 5.5 | 0.6 | 10.9 | 1.6 | 7.1 | 0.9 | 5.7 | 0.7 |
| Entries empty | 3.0 | 0.6 | 5.3 | 1.0 | 3.3 | 0.9 | 3.2 | 0.6 |
| statistics^1^ |  |  | F(1,20)=4.58 | p=0.0448 |  |  |  |  |
| statistics^2^ | F(1,20)=9.46 | p=0.006 |  |  |  |  |  |  |
| **Social novelty preference** | | | | | | | | |
| stranger 1 (s) | 113.7 | 19.2 | 172.0 | 21.3 | 138.2 | 17.0 | 220.2 | 20.0 |
| stranger 2 (s) | 288.0 | 24.3 | 243.7 | 29.6 | 262.0 | 20.9 | 174.5 | 21.6 |
| statistics | F(1,20)=20.46 | p=0.0002 |  |  | F(1,22)=11.23 | p=0.0029 |  |  |
| Entries stranger 1 | 4.1 | 1.0 | 8.4 | 1.1 | 5.8 | 1.2 | 6.9 | 0.9 |
| Entries stranger 2 | 5.6 | 1.0 | 9 | 0.9 | 5.8 | 1.1 | 6.7 | 0.7 |
| statistics^2^ | F(1,20)=11.44 | p=0.003 |  |  |  |  |  |  |
| N= | 11 |  | 11 |  | 11 |  | 13 |  |

^1^ by genotype x side

^2^ by genotype

**Table 3-1 –Morris water maze (refers to Fig. 2)**

|  | **Males** | | | | **Females** | | | |
| --- | --- | --- | --- | --- | --- | --- | --- | --- |
|  | **ctrl** | | **CKO** | | **ctrl** | | **CKO** | |
|  | **mean** | **SEM** | **mean** | **SEM** | **mean** | **SEM** | **mean** | **SEM** |
| **Visible platform test escape latency (s)** | | | | | | | | |
| Day 1 | 14 | 2 | 16 | 2 | 15 | 1 | 28 ^1^ | 4 |
| Day 2 | 7 | 1 | 8 | 1 | 9 | 1 | 11 | 1 |
| **Swim speed (cm/s)** | | | | | | | | |
| Day 1  visible platform | 16 | 0.6 | 16 | 0.8 | 16 | 1 | 15 | 1 |
| statistics^2^ |  |  |  |  |  |  | F(1,22)=9.46 | p=0.0055 |
| Day 1  hidden platform | 18 | 1 | 20 | 0.7 | 18 | 0.9 | 19 | 0.9 |
| Day 1  reversal learning | 18 | 0.6 | 17 | 1 | 19 | 1 | 20 | 0.7 |
| **Hidden platform test escape latency (s)** | | | | | | | | |
| Day 1 | 30.9 | 4.3 | 28.6 | 3.6 | 20.5 | 2.8 | 24.7 | 3.7 |
| Day 2 | 16.8 | 1.8 | 23.2 | 4.5 | 12.4 | 2.0 | 18.4 | 2.4 |
| Day 3 | 9.8 | 1.3 | 16.0 | 3.8 | 12.6 | 2.4 | 11.0 | 1.3 |
| Day 4 | 9.2 | 1.8 | 13.5 | 4.5 | 12.3 | 2.3 | 14.1 | 2.3 |
| **Hidden platform test probe trial (number of quadrant crosses)** | | | | | | | | |
| Target | 4.8 | 0.7 | 5.1 | 0.9 | 5.5 | 0.9 | 4.1 | 0.7 |
| Opposite | 2.4 | 0.4 | 2.7 | 0.9 | 1.5 | 0.3 | 3.5 | 0.5 |
| statistics^3^ | F(1,19)=6.72 | p=0.0179 |  |  | F(1,22)=10.53 | p=0.0037 |  |  |
| statistics^4^ |  |  |  |  | F(1,22)=5.67 | p=0.0264 |  |  |
| **Reversal learning test escape latency (s)** | | | | | | | | |
| Day 1 | 23.7 | 2.9 | 37.3 | 3.5 | 37.6 | 4.8 | 29.9 | 3 |
| Day 2 | 19.1 | 3.8 | 28.7 | 3.6 | 20.5 | 2.3 | 17.1 | 3.2 |
| Day 3 | 15.9 | 3.5 | 19.5 | 5.0 | 12.9 | 1. | 16.2 | 2.9 |
| Day 4 | 13.2 | 2.2 | 14.1 | 3.0 | 11.0 | 2.8 | 13.6 | 2.0 |
| **Reversal learning test probe trial (number of quadrant crosses)** | | | | | | | | |
| Target | 5.5 | 0.7 | 3.8 | 0.6 | 4.8 | 0.7 | 3.9 | 0.5 |
| Opposite | 2.4 | 0.4 | 2.9 | 0.6 | 1.9 | 0.4 | 2.4 | 0.6 |
| statistics^3^ | F(1,19)=5.09 | p=0.036 |  |  | F(1,22)=16.9 | p=0.0005 |  |  |
| statistics^5^ | F(1,19)=16.9 | p=0.0006 |  |  |  |  |  |  |
| N= | 11 |  | 10 |  | 11 |  | 13 |  |

^1^ p<0.01, comparison to female WT.

^2^ by genotype

^3^ by genotype x quadrant

^4^ by genotype x quadrant (ctrl. vs. CKO opposite quadrant)

^5^ within-genotype x quadrant ctrl.

**Table 4-1 – T maze (refers to Fig. 3)**

|  | **ctrl** | | | | | **CKO** | | | |  |  |  |  |
| --- | --- | --- | --- | --- | --- | --- | --- | --- | --- | --- | --- | --- | --- |
|  | **mean** | | | **SEM** | | **mean** | | **SEM** | |  |  |  |  |
| **Habituation distance moved (m)** | | | | | | | | | |  |  |  |  |
| Day 1 | 12.2 | | | 0.6 | | 12.1 | | 0.4 | |  |  |  |  |
| Day 2 | 10.4 | | | 0.9 | | 11.2 | | 0.5 | |  |  |  |  |
| **Training success rate** | | | | | | | | | |  |  |  |  |
| Day 1 | 0.48 | | | 0.04 | | 0.43 | | 0.04 | |  |  |  |  |
| Day 2 | 0.46 | | | 0.05 | | 0.56 | | 0.05 | |  |  |  |  |
| Day 3 | 0.58 | | | 0.05 | | 0.52 | | 0.04 | |  |  |  |  |
| Day 4 | 0.59 | | | 0.05 | | 0.65 | | 0.05 | |  |  |  |  |
| Day 5 | 0.66 | | | 0.05 | | 0.66 | | 0.05 | |  |  |  |  |
| Day 6 | 0.73 | | | 0.05 | | 0.66 | | 0.05 | |  |  |  |  |
| Day 7 | 0.82 | | | 0.04 | | 0.76 | | 0.05 | |  |  |  |  |
| Day 8 | 0.86 | | | 0.03 | | 0.73 | | 0.06 | |  |  |  |  |
| Day 9 | 0.93 | | | 0.03 | | 0.80 | | 0.04 | |  |  |  |  |
| Day 10 | 0.94 | | | 0.02 | | 0.82 | | 0.05 | |  |  |  |  |
| Day 11 | 0.92 | | | 0.03 | | 0.88 | | 0.04 | |  |  |  |  |
| Day 12 | 0.98 | | | 0.01 | | 0.92 | | 0.04 | |  |  |  |  |
| Day 13 | 0.95 | | | 0.02 | | 0.94 | | 0.03 | |  |  |  |  |
| Day 14 | 0.96 | | | 0.02 | | 0.91 | | 0.03 | |  |  |  |  |
| **Probe trial strategy (absolute)** | | | | | | | | | |  |  |  |  |
|  | P | | | R | | P | | R | |  |  |  |  |
| Probe trial 1 | 10 | | | 23 | | 14 | | 17 | |  |  |  |  |
| Probe trial 2 | 9 | | | 24 | | 13 | | 18 | |  |  |  |  |
| **Probe trial strategy (relative)** | | | | | | | | | |  |  |  |  |
|  | P | | | R | | P | | R | |  |  |  |  |
| Probe trial 1 | 0.30 | | | 0.70 | | 0.45 | | 0.55 | |  |  |  |  |
| statistics | ctrl. | | |  | | p=0.303 | | vs. ctrl | |  |  |  |  |
| Probe trial 2 | 0.27 | | | 0.73 | | 0.42 | | 0.58 | |  |  |  |  |
| statistics | ctrl. | | |  | | p=0.294 | | vs. ctrl | |  |  |  |  |
| N= | 33 | | |  | | 31 | |  | |  |  |  |  |
| **Strategy transitions probe trials** | | | | | | | | | | | | | |
|  | | **ctrl** | | | | | | | **CKO** | | | | |
|  | | R🡪P | P | | R | | P🡪R | | R🡪P | | P | R | P🡪R |
| Absolute | | 5 | 5 | | 17 | | 6 | | 9 | | 4 | 8 | 10 |
| Relative | | 0.15 | 0.15 | | 0.52 | | 0.18 | | 0.29 | | 0.13 | 0.26 | 0.32 |
| statistics | |  |  | | ctrl. | |  | |  | |  | p=0.0434 | vs. ctrl |

**Table 5-1 – CognitionWall (refers to Fig. 4)**

|  | **Males** | | | | **Females** | | | | |
| --- | --- | --- | --- | --- | --- | --- | --- | --- | --- |
|  | **ctrl** | | **CKO** | | **ctrl** | | | **CKO** | |
|  | **mean** | **SEM** | **mean** | **SEM** | **mean** | | **SEM** | **mean** | **SEM** |
| **Total distance moved (m)** | | | | | | | | | |
| Day 1-4 | 1908 | 181 | 2396 | 302 | 2244 | | 119 | 2383 | 134 |
| statistics | ctrl. |  | p=0.0409 | vs. m ctrl | p=0.0106 | | vs. m ctrl | p=0.2959 | vs. f ctrl |
| **Total wall entries** | | | | | | | | | |
| Day 1-4 | 2887 | 318 | 2717 | 237 | 3222 | | 139 | 3199 | 140 |
| statistics | ctrl. |  | p=0.4475 | vs. m ctrl | p=0.0731 | | vs. m ctrl | p=0.8567 | vs. f ctrl |
| **Errors to 80% learning criterion** | | | | | | | | | |
| Day 1-2 | 138 | 27 | 119 | 17 | 156 | | 49 | 170 | 17 |
| statistics | ctrl. |  | p=0.7664 | vs. m ctrl | p=0.7984 | | vs. m ctrl | p=0.8777 | vs. f ctrl |
| Day 3-4 | 494 | 73 | 444 | 40 | 448 | | 37 | 474 | 106 |
| statistics | ctrl. |  | p=04321 | vs. m ctrl | p=0.5151 | | vs. m ctrl | p=0.7803 | vs. f ctrl |
| **Rewards earned per day** | | | | | | | | | |
| Day 1 | 72 | 18 | 75 | 13 | 94 | 11 | | 101 | 7 |
| statistics | ctrl. |  | p=0.9001 | vs. m ctrl | p=0.2849 | vs. m ctrl | | p=0.6645 | vs. f ctrl |
| Day 2 | 112 | 9 | 113 | 8 | 115 | 5 | | 125 | 8 |
| statistics | ctrl. |  | p=0.9559 | vs. m ctrl | p=0.8621 | vs. m ctrl | | p=0.5717 | vs. f ctrl |
| Day 3 | 86 | 26 | 58 | 18 | 88 | 14 | | 93 | 20 |
| statistics | ctrl. |  | p=0.2739 | vs. m ctrl | p=0.9129 | vs. m ctrl | | p=0.7896 | vs. f ctrl |
| Day 4 | 117 | 20 | 93 | 21 | 131 | 8 | | 107 | 12 |
| statistics | ctrl. |  | p=0.3469 | vs. m ctrl | p=0.4728 | vs. m ctrl | | p=0.1564 | vs. f ctrl |

**Table 6-1 – Fear conditioning (refers to Fig. 5)**

|  | **ctrl** | | **CKO** | |
| --- | --- | --- | --- | --- |
|  | **mean** | **SEM** | **mean** | **SEM** |
| **Acquisition** | | | | |
| BL (%) | 0.5 | 0.2 | 0.6 | 0.2 |
| Tone 1 (%) | 1.6 | 1.4 | 2.4 | 1.0 |
| Tone 2 (%) | 5.9 | 2.5 | 11.8 | 3.0 |
| Tone 3 (%) | 46.5 | 8.4 | 57.2 | 7.8 |
| statistics | Sex: F(1,35) = 0.054 | p = 0.818 | Genotype: F(1,35) = 3.230 | p = 0.081 |
| **Extinction day 1** | | | | |
| BL (%) | 32.9 | 6.0 | 31.9 | 5.8 |
| Tone 1-5 (%) | 61.4 | 7.6 | 60.9 | 6.3 |
| Tone 6-10 (%) | 58.4 | 8.1 | 59.9 | 6.2 |
| Tone 11-15 (%) | 52.2 | 7.3 | 52.3 | 6.6 |
| Tone 16-20 (%) | 50.4 | 8.0 | 49.5 | 7.0 |
| statistics | Sex: F(1,35) = 0.034 | p = 0.854 | Genotype: F(1,35) = 0.008 | p = 0.931 |
| **Extinction day 2** | | | | |
| BL (%) | 32.9 | 6.3 | 31.9 | 5.6 |
| Tone 1-5 (%) | 60.2 | 7.2 | 60.3 | 6.6 |
| Tone 6-10 (%) | 40.6 | 7.0 | 49.1 | 6.9 |
| Tone 11-15 (%) | 47.1 | 7.4 | 49.0 | 6.5 |
| Tone 16-20 (%) | 36.2 | 7.4 | 49.0 | 7.0 |
| statistics | Sex: F(1,35) = 0.033 | p = 0.858 | Genotype: F(1,35) = 0.295 | p = 0.591 |
| **Extinction day 3** | | | | |
| BL (%) | 12.3 | 3.0 | 20.5 | 3.9 |
| Tone 1-5 (%) | 46.5 | 6.7 | 48.9 | 5.5 |
| Tone 6-10 (%) | 39.4 | 6.4 | 43.5 | 5.2 |
| Tone 11-15 (%) | 36.6 | 6.0 | 40.5 | 6.9 |
| Tone 16-20 (%) | 27.3 | 5.5 | 38.8 | 5.1 |
| statistics | Sex: F(1,35) = 1.442 | p = 0.238 | Genotype: F(1,35) = 0.993 | p = 0.326 |
| **Renewal** | | | | |
| BL (%) | 34.5 | 6.1 | 39.1 | 5.4 |
| Tone 1 (%) | 49.0 | 8.0 | 74.5 | 6.1 |
| statistics | Sex: F(1,35) = 1.137 | p = 0.294 | Genotype: F(1,35) = 5.293 | p = 0.027 |
| Tone 2 (%) | 49.1 | 7.7 | 64.1 | 7.2 |
| Tone 3 (%) | 53.4 | 7.9 | 64.1 | 6.5 |
| N= | 18 |  | 21 |  |
